# Hypothalamic effective connectivity at rest is associated with body weight and energy homeostasis

**DOI:** 10.1101/2021.06.22.449523

**Authors:** Katharina Voigt, Zane B. Andrews, Ian H. Harding, Adeel Razi, Antonio Verdejo-Garcia

## Abstract

Hunger and satiety states drive eating behaviours via changes in brain function. The hypothalamus is a central component of the brain networks that regulate food intake. Animal research parsed the roles of the lateral hypothalamus (LH) and the medial hypothalamus (MH) in hunger and satiety respectively. Here, we examined how hunger and satiety change information flow between human LH and MH brain networks, and how these interactions are influenced by body mass index. Forty participants (15 overweight/obese) underwent two resting-state functional MRI scans: after overnight fasting (fasted state) and following a standardised meal (sated state). The direction and valence (excitatory/inhibitory influence) of information flow between the MH and LH was modelled using spectral dynamic causal modelling. Our results revealed two core networks interacting across homeostatic state and weight status: subcortical bidirectional connections between the LH, MH and the substantia nigra pars compacta (prSN), and cortical top-down inhibition from frontoparietal and temporal areas. During fasting relative to satiety, we found higher inhibition between the LH and prSN, whereas the prSN received greater top-down inhibition from across the cortex. Individuals with higher BMI showed that these network dynamics occur irrespective of fasted or satiety states. Our findings reveal fasting affects brain dynamics over a distributed hypothalamic-midbrain-cortical network. This network is less sensitive to state-related fluctuations among people with obesity.

## Introduction

The hypothalamus accounts for only approximately 3% of total human brain tissue, but is one of the most vital structures regulating a plethora of bodily functions essential for survival (Stuber & Wise, 2016). This small subcortical region regulates our response to stress, arousal, reward processing, body temperature, fertility and sexual behaviour, motivation and food intake (Pop et al., 2018). Early preclinical lesion studies subdivided the hypothalamus anatomically and functionally into lateral hypothalamus (LH) and medial hypothalamus (MH), leading to the concept of a “dual centre model” (Anand & Brobeck, 1951; Brobeck et al., 1943; Elmquist et al., 1999; Hetherington & Ranson, 1983). Lesions to the MH resulted in increased appetite, food intake and weight gain, marking the MH as “satiety centre”. Lesions to the LH in turn induced abnormal decreases in appetite and food intake, labelling the LH as “hunger centre”. More recent studies show that neurons in the LH regulate food consumption and appetitive motivation with extensive reciprocal connections to the dopaminergic midbrain governing reward processing in support of goal-directed food seeking (Jennings et al., 2013; Jennings et al., 2015; Nieh et al., 2016; Rossi & Stuber, 2018). The LH is a large single region with numerous heterogeneous neuronal populations, whereas the MH can be further subdivided into many important nuclei involved in the regulation of food intake, blood glucose and weight control. This includes the arcuate nucleus, the ventromedial hypothalamic nucleus, the dorsomedial hypothalamic nucleus and the paraventricular nucleus. Both the LH and MH nuclei function in a metabolic state-dependent manner and can be reshaped by obesity and energy homeostasis (Chen et al., 2017; Rossi et al., 2019). Moreover, these hypothalamic areas are heavily integrated into intra- and inter-hypothalamic neural circuits and networks, with the majority of LH connectivity coming from outside the LH (Burdakov & Karnani, 2020).

Although animal research has greatly contributed to understanding how hypothalamic neural circuits integrate peripheral and central signals to control food intake, the connectivity in humans to and from the LH and MH nuclei remains poorly understood. Energy homeostasis relies on the coordinated and dynamic interactions of the hypothalamus both to (bottom-up) and from (top-down) a broad set of cortical and subcortical brain regions (Rossi & Stuber, 2018). A precise description of how the LH and MH network functions in response to changes in homeostatic state in humans is thus required to bridge the gap between animal and human research, and to provide a critical step towards defining the neural underpinnings of maladaptive eating patterns leading to obesity in humans. An examination of the LH and MH networks is further supported by the well-known psychological comorbidities associated with metabolic diseases such as anorexia, obesity and diabetes (Florent et al., 2020; Penninx & Lange, 2018).

Research has begun to establish the links between the hypothalamic network and obesity using functional magnetic resonance imaging (fMRI). One study by Kullmann and colleagues (2014) described differences in the hypothalamic network between people with excess weight and those with healthy weight. Functional connectivity analyses revealed the LH was more heavily connected to the dorsal striatum, anterior cingulum, and frontal operculum, and the MH was more connected to the medial orbitofrontal cortex and nucleus accumbens (replicated recently by Zhang et al., 2018). Further, in participants with excess weight, the functional connectivity of the MH, but not the LH, was increased with the nucleus accumbens and medial prefrontal cortex. These results highlight the existence of two distinct circuitries originating from the MH and LH that are modulated by obesity. However, these studies do not reveal the functional interactions (e.g., inhibition or excitation) nor do they differentiate between bottom-up and top-down effects within the network. Further, given the metabolic state-dependency of the LH and MH circuitries (Chen et al., 2017; Rossi et al., 2019), it is also critical to investigate this network as a function of more dynamic state-dependent changes in energy homeostasis, such as in states of fasting vs. satiety.

The current study examines the directionality (bottom-up vs. top-down) and valence (inhibition vs. excitation) of connections of the LH and MH with key cortical and subcortical brain regions. These network dynamics are examined in participants varying in weight (healthy vs. excess weight) in a fasted or sated state. We capitalise on recent advances in modelling the interactions within a brain network based on the low-frequency endogenous fluctuations in resting-state functional magnetic resonance imaging (rsfMRI) data using spectral dynamic causal modelling (spDCM; Friston et al. 2014; Razi et al. 2015; Park et al. 2018) and state of the art anatomical labelling (Rolls et al., 2020). In contrast to conventional functional connectivity analyses (e.g., Kullmann et al., 2014; Zhang et al., 2018), spDCM predicts directional communications among distributed brain regions (i.e., effective connectivity; Friston, Harrison, & Penny, 2003). We hypothesise that (i) the LH and MH would show distinct functional connectivity and (ii) homeostatic state and weight would change effective connectivity within LH and MH networks.

## Methods

### Participants

Forty participants were recruited via flyers and social-media advertisements. Participants were required to be 18 – 55 years old, right-handed, and have a Body Mass Index (BMI) between 18 and 30kg/m^2^. Screening criteria excluded people with a history of hypertension or diabetes, neurological or psychiatric illness, or who had recently taken psychoactive medications. Additionally, participants could not be subject to MRI contradiction, such as metal implants or pregnancy. The number of participants was chosen based on a sample size estimation study revealing that 20 participants provided for reliable DCM predictions (Goulden et al., 2012). In agreement, recent research showed robust model predictions using similar sample sizes when applying spDCM to rsfMRI data (Park et al., 2018; Preller et al., 2019; Voigt et al., 2020). Out of the 40 participants, two were excluded from analysis as they did not complete both fasted and sated rsfMRI scans. In total, data from 38 participants were included into the analyses (Table 1 for participants’ demographics). All participants gave written consent before participating and were reimbursed with $100 gift card vouchers. The Monash University Human Research Ethics Committee approved the study (2019-5979-30222) following the Declaration of Helsinki.

**Table 1.**
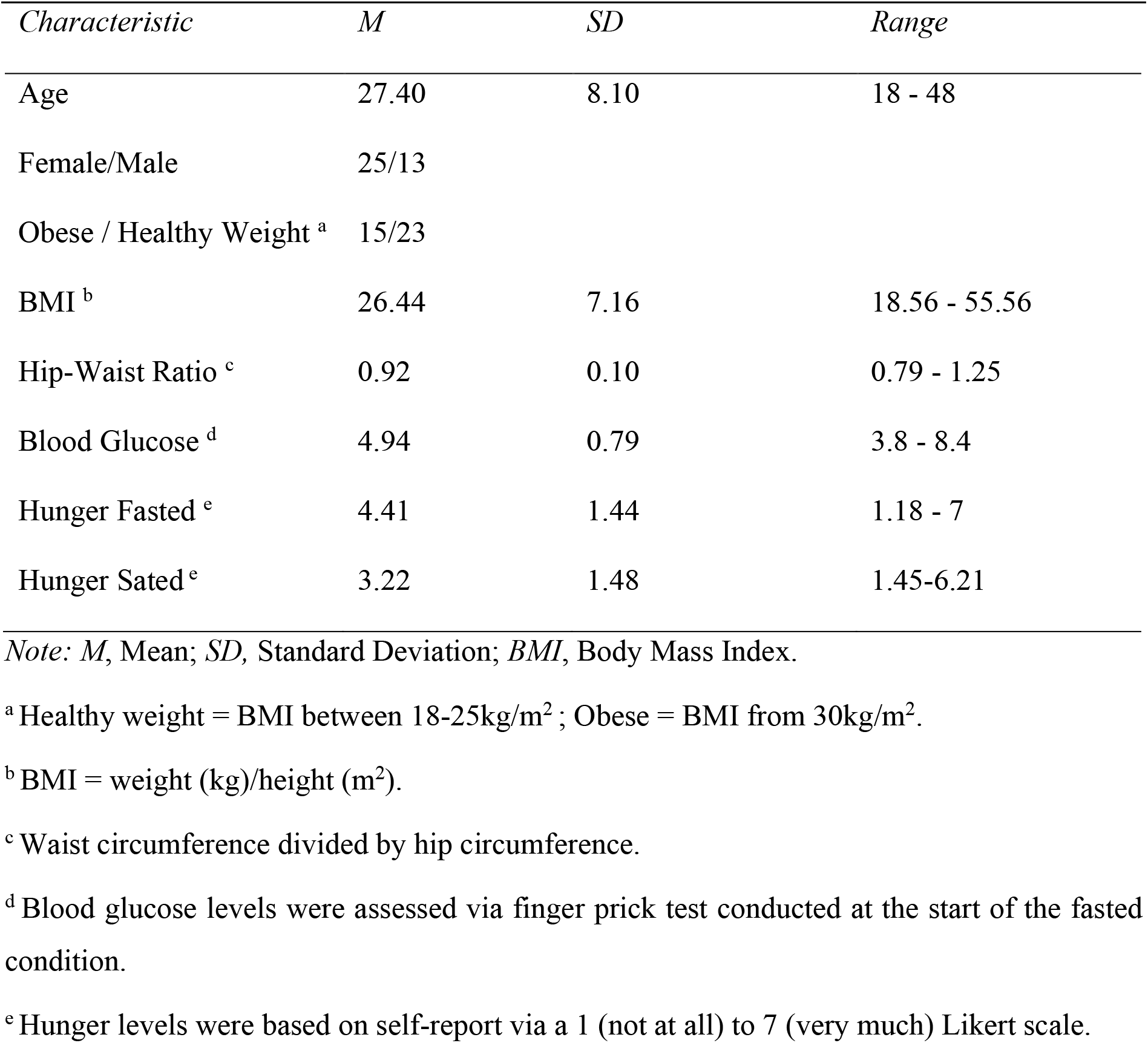
Participants’ demographics.

### Experimental procedure

Participants completed two resting-state fMRI scans, one after an overnight fast (fasted condition) and one after a standard breakfast (sated condition). In both conditions, participants were instructed to have a standard meal (700-1000kJ) between 7.30pm and 8.30pm on the night prior to their scan and subsequently to refrain from eating or drinking (except for water) until their morning scan. Fasting blood glucose levels were measured via a standard finger prick test. For the sated condition, participants received a breakfast (293 kcal) one hour prior to their scan. Subjective self-reports of hunger (1 = not hungry at all; 7 = very hungry) revealed a significant difference in the perception of hunger during the fasted (*M* = 4.42; *SD* = 1.44) and sated (*M* = 3.22; *SD* = 1.48) condition (*t*(36) = 4.72, *p* < 0.001). There was no interaction between subjective reported hunger and BMI, but there was between BMI and fasting blood glucose level (see Appendix, Figure S1 and Figure S2). All scans were scheduled in the morning between 9am and 10am. On average there were 5.82 days (*SD* = 3.73 days) between the two scanning sessions. The order of fasted and sated scans was counterbalanced across participants.

### Resting-state fMRI data acquisition

Resting-state fMRI data were acquired using a 3-Tesla Siemens Skyra MRI scanner equipped with a 32-channel head coil at the Monash Biomedical Imaging Research Centre (Melbourne, Victoria, Australia). During a total acquisition time of 7.8 minutes, 600 volumes were acquired for each participant and homeostatic condition using a multiband gradient echo pulse sequence (45 axial slices; time of repetition, TR = 780 ms; echo time, TE = 21 ms, resolution 3×3×3mm). In order to obtain structural brain information for each participant, a high-resolution T1-weighted magnetisation-prepared rapid gradient echo (MPRAGE) covering the whole brain was measured (repetition time = 2300ms; echo time = 2.07 ms; flip angle = 9°; 192 slices; field of view = 256×256 mm, voxel resolution = 1mm isotropic). Participants were instructed to rest while fixating on a central black crosshair (i.e. eyes-open resting-state protocol).

Following the resting-state fMRI scan of each session, participants completed a task-based fMRI session, during which they were required to make realistic choices between unhealthy and healthy drinks. The full study protocol and task-based results are reported elsewhere (Harding et al., 2018). The task-based fMRI session was always performed after the resting-state MRI session, to avoid confounding the resting-state BOLD signals (Tambini et al., 2009).

### Resting-state fMRI data analyses

#### Preprocessing

Functional images were preprocessed using SPM12 (revision 12.2, www.fil.ion.ucl.ac.uk). The preprocessing steps consisted of spatial realignment, tissue segmentation and spatial normalisation to the standard EPI template of the Montreal Neurological Institute (MNI), and spatial smoothing using a Gaussian kernel of 6-mm FWHM. None of the participants exceeded excessive head motion of larger than 3mm. For the seed-based functional connectivity analyses, we applied an additional temporal bandpass filter (0.01-0.08Hz) to remove low frequency drifts and high frequency physiological noise as well as linear detrending the data. Nuisance covariate regression was performed to remove signal variance of non-neuronal origin using timeseries extracted from the white matter, and independently from the cerebrospinal fluid, in addition to the six parameters define the magnitude of frame-by-frame head motion (3 x translation; 3 x rotation).

### Statistical fMRI Analyses

We first conducted an initial seed-based functional connectivity analysis (using the bilateral MH and LH as seeds, Figure 1). This analysis was used to obtain the brain areas that are associated with the MH and LH at rest (i.e., the hypothalamic functional resting state network). Next, we conducted a spDCM analyses to investigate the causal interactions between these areas and how they differ as a function of homeostatic state (fasted vs. sated), BMI and the interaction between homeostatic state and BMI. The details of these two analyses are outlined next.

**Figure 1.**
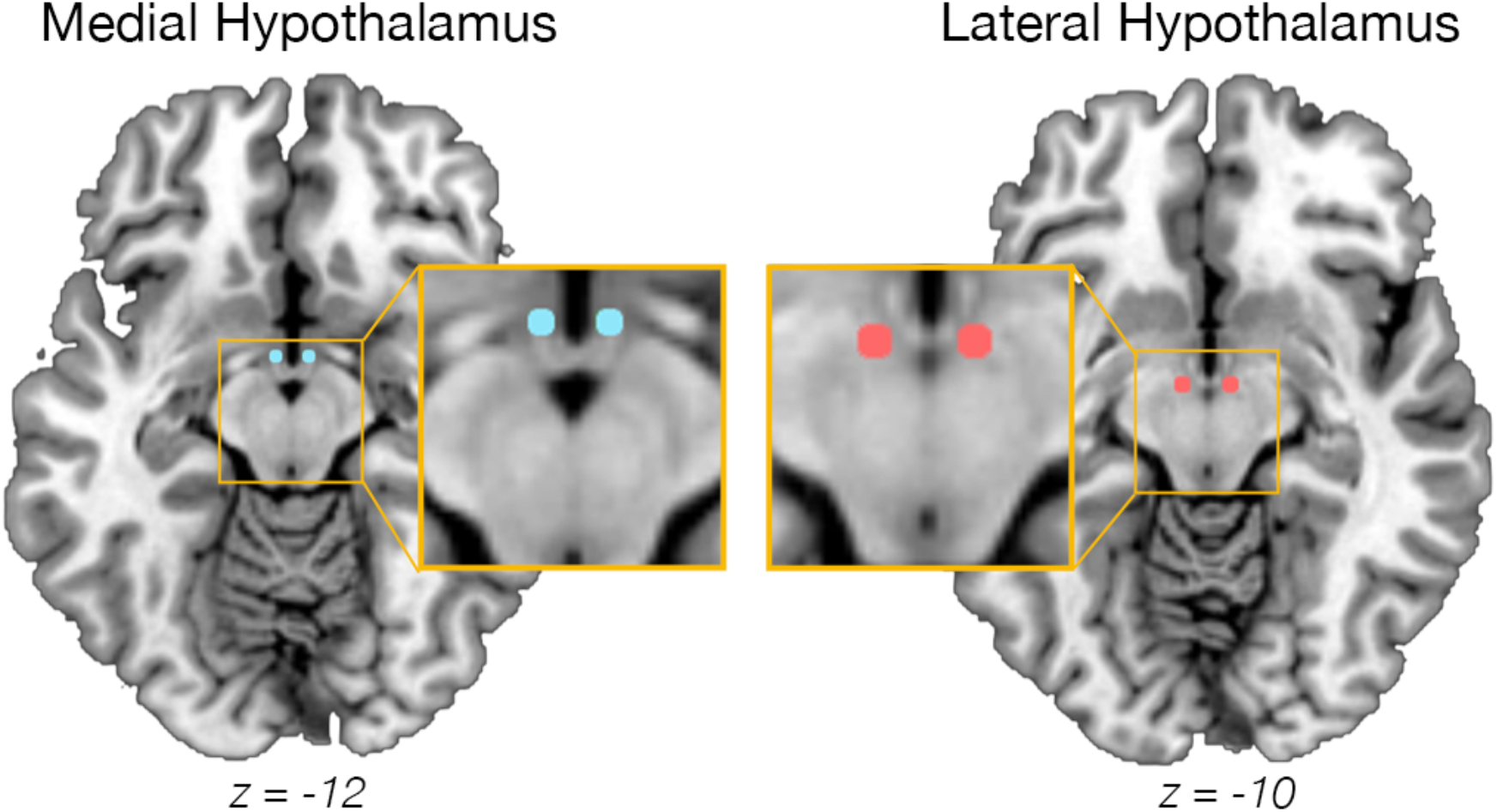
Seed regions of the medial hypothalamus (MH; MNI coordinates x = ±4, y = -2, z = -12), and lateral hypothalamus (LH; MNI coordinates x= ±6, y= -9, -10) (based on Baroncini et al., 2012) used to obtain the hypothalamic network for subsequent spDCM analysis.

### Seed-based functional connectivity analysis

Functional connectivity (FC) maps of the hypothalamic functional resting state network were obtained using an initial seed-based functional connectivity analyses across all subjects. We defined two ROIs according to (Baroncini et al., 2012): the bilateral lateral hypothalamus (LH; MNI coordinates x = ± 6, y = -9, z = -10) and bilateral medial hypothalamus (MH; MNI coordinates x = ± 4, y = -2, z = -12) using 2-mm-radius spheres (Figure 1). To minimise overlap between the two ROIs, we chose the peak voxel of the LH to be in the posterior part of the LH according to Baroncini and collegues (2012). The seeds were, as such, spatially separated by > 6mm (i.e., > 1mm after smoothing).

In order to define the general hypothalamus network that was associated with either of the LH or MH across subjects, we extracted the average timeseries from LH and MH combined. This time series was then correlated with the timeseries of activity within each of voxel across the rest of the brain. The resulting FC maps were transferred to z-scores using Fisher’s transformation and analysed using a one-sample’s t-test in SPM12 (Wellcome Department of Cognitive Neurology, London, UK). Brain voxels with a threshold of p < 0.05, family-wise error (FWE) corrected for multiple comparisons on the voxel-level were considered significant. Anatomical regions were labelled using the automatic anatomical labelling atlas (AAL3, Rolls et al., 2020). This atlas includes brain areas that have not generally been defined in other atlases, such as subdivisions of the thalamus. Previous studies investigating the functional connectivity of the MH and LH (Kullmann et al., 2014; Zhang et al., 2018) have not used such precise anatomical labelling.

### Spectral Dynamic Causal Modeling

The spDCM analyses were performed using the functions of DCM12 (revision 7196) implemented in SPM12 (version 7487) in Matlab 2018b. In order to address our main hypotheses, we focused on spDCM analyses that assessed (1) effective connectivity of the hypothalamic network in the fasted and sated states independently, (2) changes in hypothalamic effective connectivity between the fasted versus sated state, independent of BMI (main effect of fasting), (3) changes in hypothalamic effective connectivity modulated by BMI, independent of energy state (main effect of BMI), and (4) changes in fasting-related effective connectivity of the hypothalamus modulated by BMI (fasting-by-BMI interaction).

#### First Level spDCM Analysis

In order to assess the effective connectivity of the hypothalamus network, regions revealed by the initial functional connectivity analyses of both the MH and LH in conjunction with a minimum voxel size of 20 were used as ROIs for the subsequent spDCM analyses (Figure 2, Table 2). At the first-level, a fully-connected model was created for each participant and each session. Next, we inverted (i.e. estimated) the DCMs using spectral DCM, which fits the complex cross-spectral density using a parameterised power-law model of endogenous neural fluctuations (Razi et al., 2015). This analysis provides measures of causal interactions between regions, as well as the amplitude and exponent of endogenous neural fluctuations within each region (Razi et al., 2015). Model inversion was based on standard variational Laplace procedures (Friston et al., 2007). This Bayesian inference method uses Free Energy as a proxy for (log) model evidence, while optimising the posterior density under Laplace approximation.

**Table 2.**
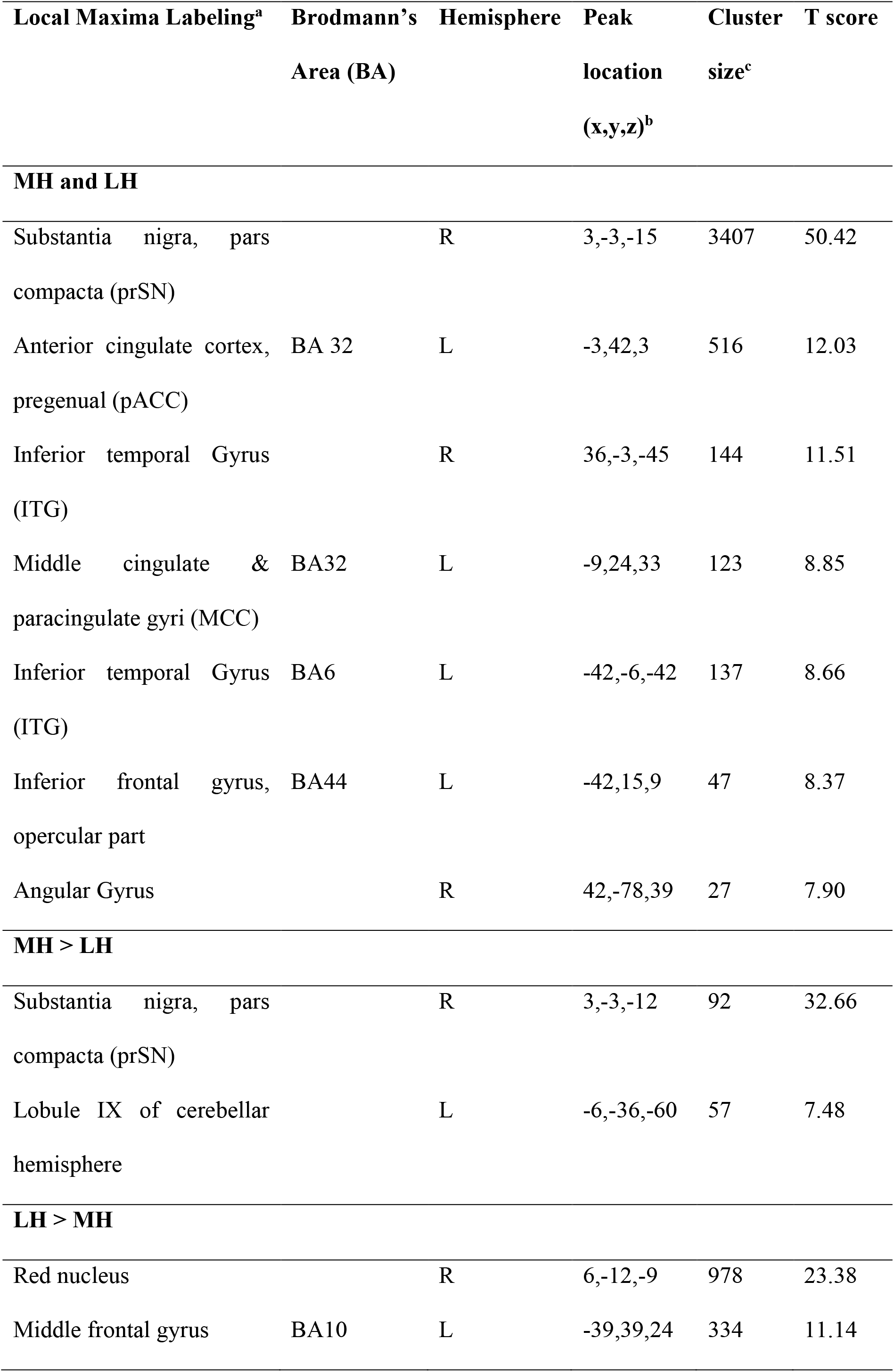
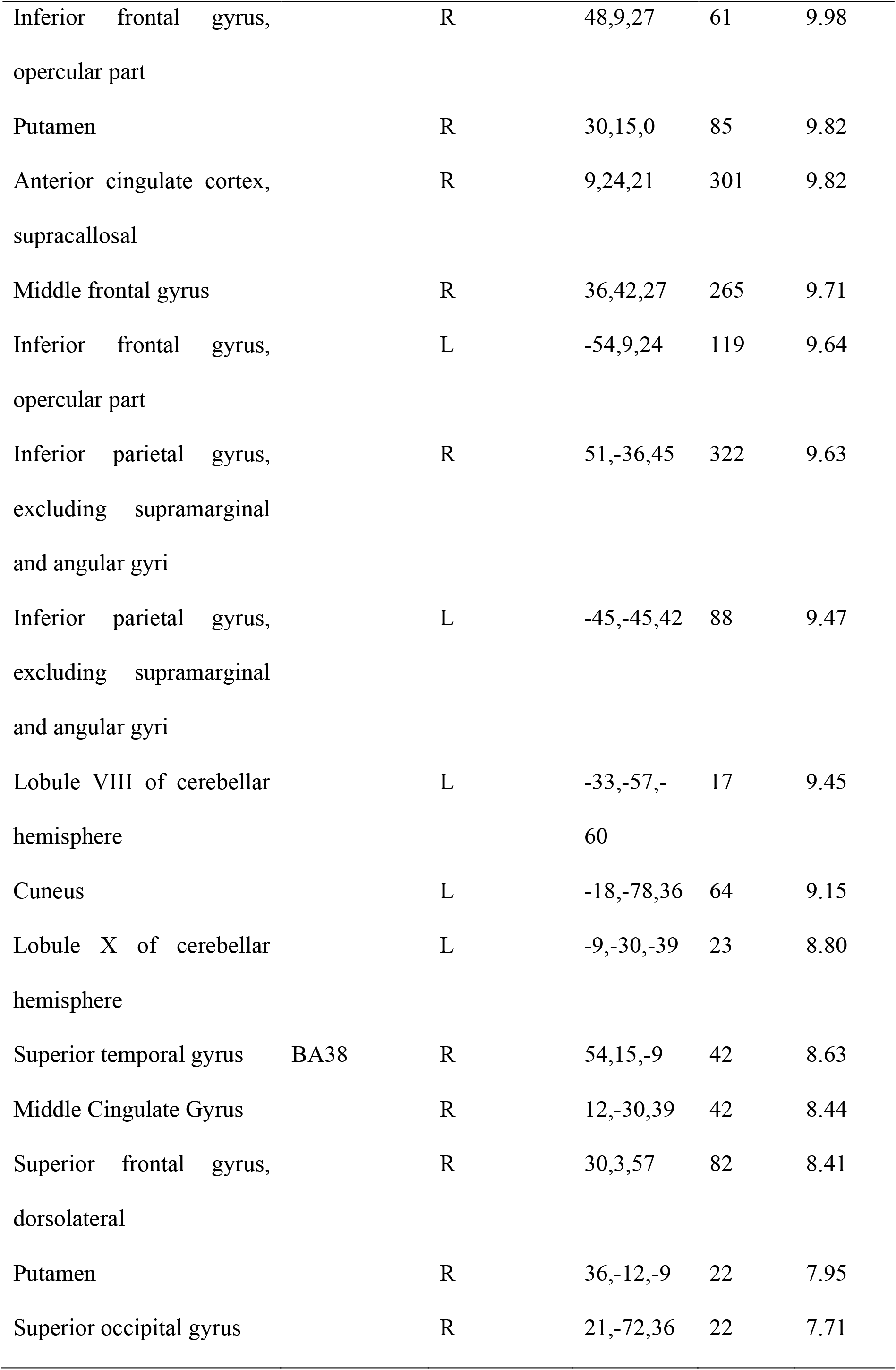
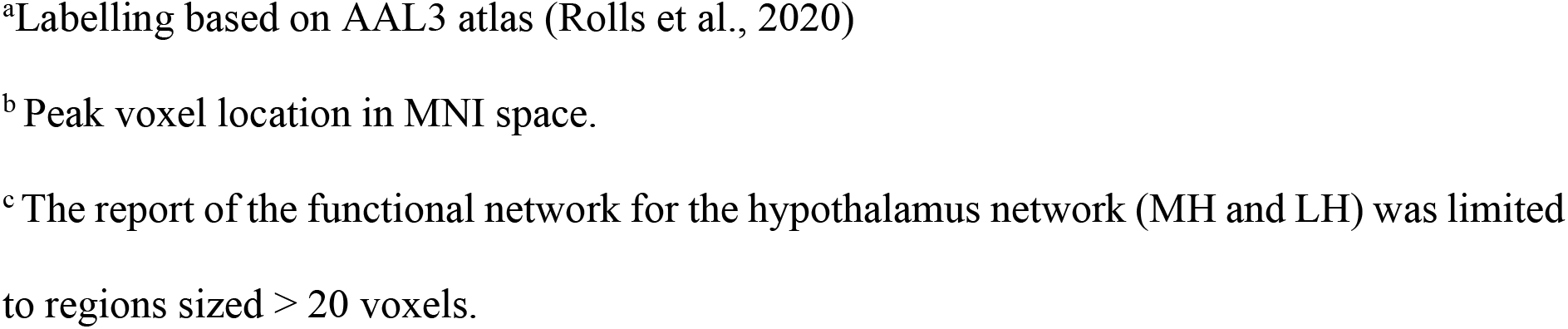
Peak coordinates of hypothalamus intrinsic functional connectivity networks

**Figure 2.**
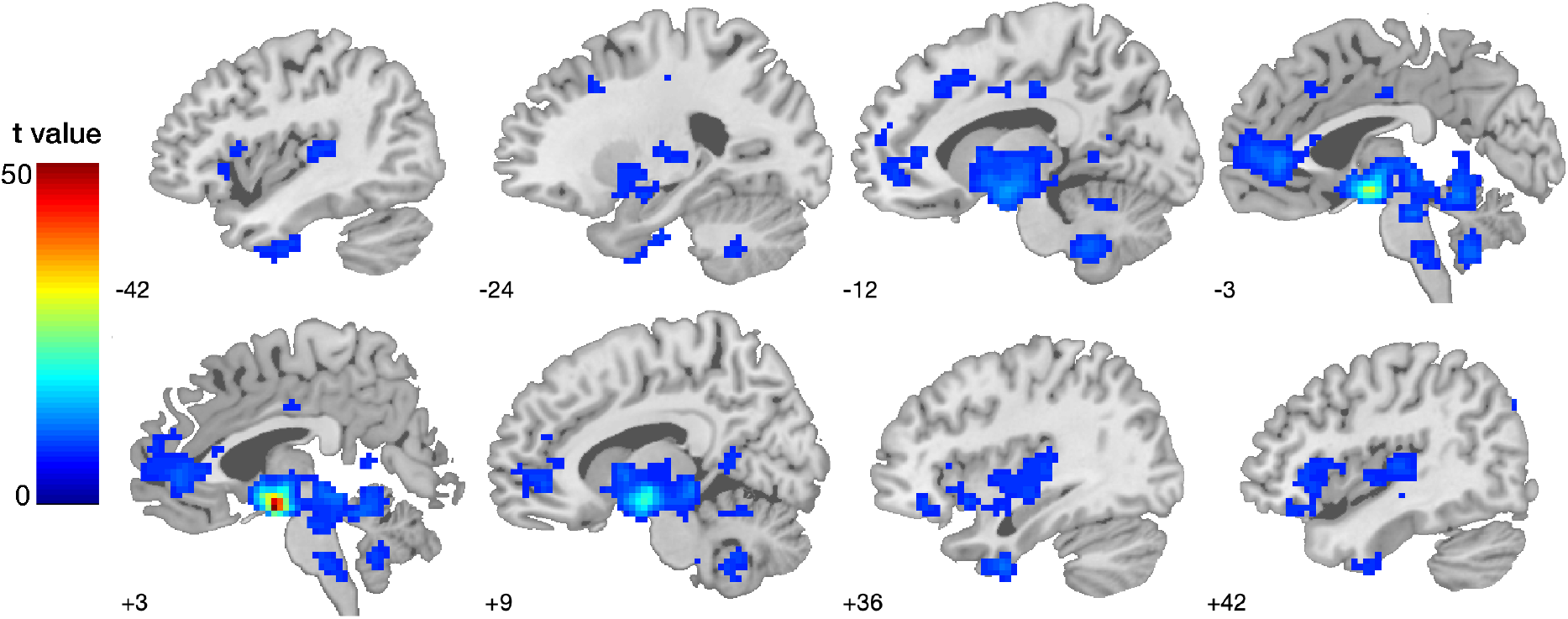
Functional connectivity network of the lateral and medial hypothalamus. Seed-based functional connectivity results by using the bilateral seed mask LH and MH (Figure 1). Results reflect whole-brain one-sample t tests at p < 0.001 FWE-corrected. For differences in MH and LH functional connectivity, refer to Table 2. The color bar represents voxel T values.

#### Second Level spDCM Analysis

To characterise how group differences in neural circuitry were modulated by BMI and energy state, hierarchical models over the parameters were specified within a hierarchical Parametric Empirical Bayes (PEB) framework for DCM (Friston et al., 2016). The four models we used were based on our hypotheses as follows. Firstly, to investigate the effective connectivity of the MH and LH, initial intercept models were estimated within the PEB framework. The intercept models provide the baseline effective connectivity in independence of any behavioural measures. Secondly, we were interested in the group difference between fasted versus sated conditions and in this PEB analysis, we contrasted the DCMs for the fasted against the sated condition whilst controlling for BMI. Thirdly, we were interested associating effective connectivity with BMI and used BMI as a main regressor of interest whilst controlling for homeostatic condition. Lastly, we were interested in interaction between group factor (fasted versus sated) and BMI and in this PEB analysis, we used the interaction between BMI and the group factor (fasted versus sated) as main variables of interest.

For each of the presented models, all behavioural regressors were mean-centered so that the intercept of each model was interpretable as the mean connectivity. We tested the relationships between all covariates (i.e., age, gender, BMI, subjective hunger reports, blood glucose levels; Supplementary Analyses), and included blood glucose as a mean-centred covariate as it correlated with BMI in interaction with the experimental condition.

Bayesian model reduction was used to test all combinations of parameters (i.e., reduced models) within each parent PEB model (assuming that a different combination of connections could exist [Friston et al., 2016]) and ‘pruning’ redundant model parameters. Parameters of the set of best-fit pruned models (in the last Occam’s window) were averaged and weighted by their evidence (i.e., Bayesian Model Averaging) to generate final estimates of connection parameters. To identify important effects (i.e., changes in directed connectivity), we compared models, using log Bayesian model evidence to ensure the optimal balance between model complexity and accuracy, with and without each effect and calculated the posterior probability for each model as a softmax function of the log Bayes factor. We treat effects (i.e., connection strengths and their changes) with a strong posterior probability > 0.99 (equivalent of very strong evidence in classical inference) as significant for reporting purposes.

Finally, in order to determine the predictive validity (e.g. whether BMI can be predicted from the final, reduced spDCM’s individual connections), leave-one-out cross validation was performed within the PEB framework (Zeidman et al., 2019). This procedure fits the PEB model in all but one participant and predicts the covariate of interest (e.g., homeostatic state) for the left-out participant. This is repeated with each participant to assess the averaged prediction accuracy for each model.

## Results

We first provide the overview of the seed-based functional connectivity analysis, which was conducted for the derivation of the hypothalamic network. Secondly, we describe the causal dynamics within this derived hypothalamic network.

### Seed-based functional connectivity analyses

The seedbased functional connectivity analyses (combined LH and MH seed mask, Figure 1) revealed a hypothalamic network comprising of seven regions in the substantia nigra, anterior and middle cingulate, inferior temporal and frontal gyrus, and the angular gyrus (Table 2). These seed-based functional connectivity results subtending the MH and LH functional connectivity were used to define the network for each participant for the subsequent effective connectivity analysis. The differential contrast (MH > LH) revealed a stronger functional connectivity between the MH and substantia nigra, cerebellum and precuneus. The lateral hypothalamus (LH > MH contrast) showed stronger functional connectivity to the red nucleus, inferior and middle frontal gyrus, putamen, anterior and middle cingulate cortex, putamen and regions of the parietal, occipital lobe as well as cerebellum, (p < 0.001, FWE-corrected, Table 2, Figure 2).

### Spectral Dynamic Causal Modelling Results

The average variance explained across subject-level DCM inversion was very high (Hunger: M = 86.78, SD = 3.23, range = 80.20-94.90; Satiety: M = 86.66, SD = 2.58, range = 80.02-91.68), indicating very good model convergence.

### Effective connectivity of the MH and LH during the Fasted and Sated Condition

Figure 3 illustrates the effective connectivity results of the LH and MH during the fasted and sated condition separately. Across the fasted and sated condition, there were decreased connections from anterior and mid-cingulate, frontal, temporal and parietal cortex to the both the LH and MH. There was no evidence for connections going from the LH or MH to these cortical regions in the fasted condition. During the sated condition, there was an increased connection from the left MH to the angular gyrus and from the right LH to the pACC.

**Figure 3.**
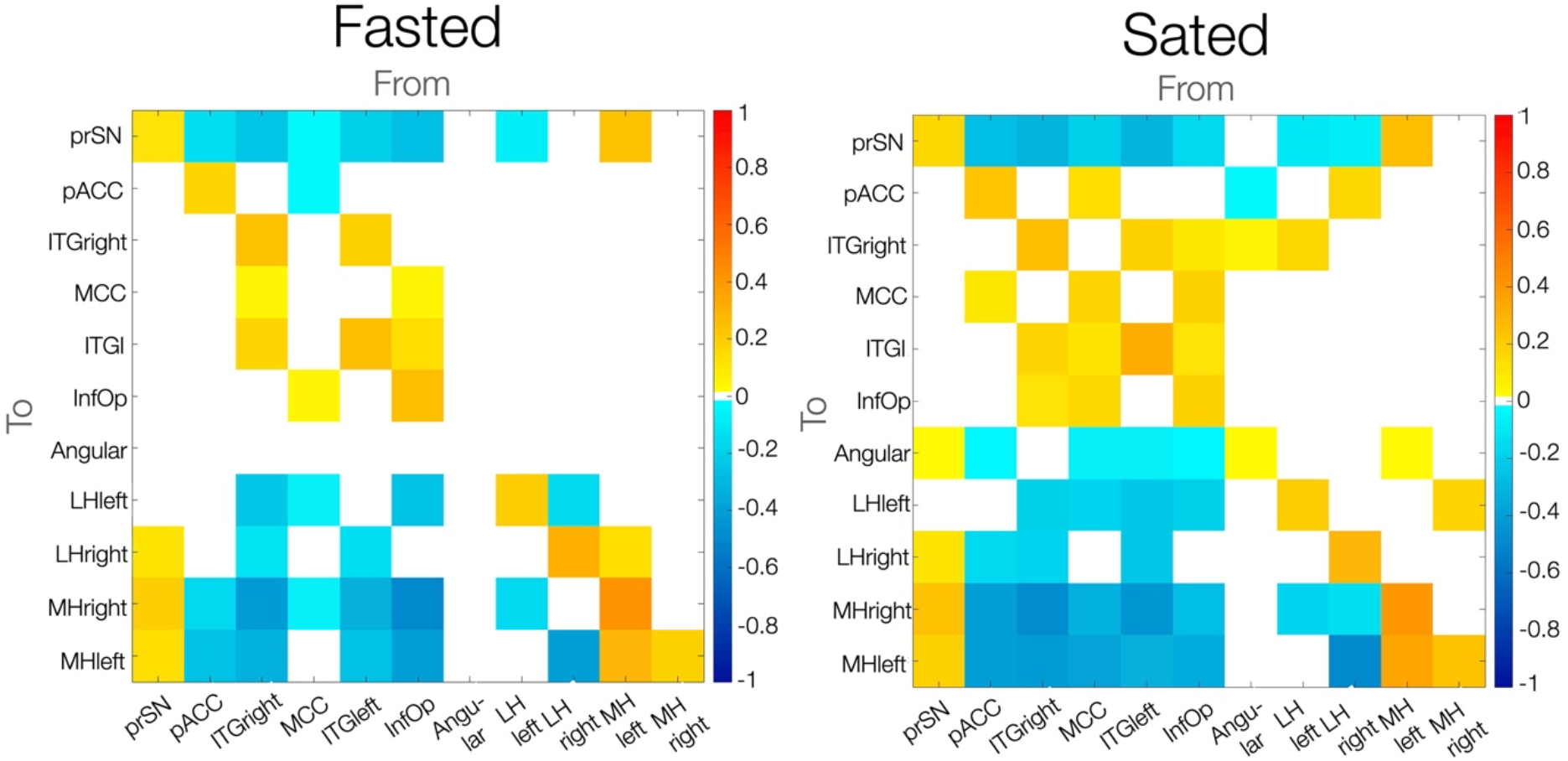
Effective resting-state connectivity of the hypothalamic network across during the fasted vs. sated condition. Colour bar indicates effect sizes in Hz. Posterior probability of > .99 (very strong evidence). Abbreviations: prSN, substantia nigra pars compacta; MH, medial hypothalamus; LH, lateral hypothalamus; pACC, anterior cingulate cortex pregenual; MCC, middle cingulate and paracingulate gyri; ITG, inferior temporal gyrus; InfOp, inferior frontal gyrus, opercular part.

### Effective connectivity of the MH and LH in fasted vs. sated states

Fasting, compared to satiety, was associated with a decreased excitatory influence from the substantia nigra to the left MH (0.01 Hz, 95% CI [-0.03, 0.004]). The left MH in turn showed an increased excitation to the left LH (0.06 Hz, 95% CI [-0.002, 0.11]) and to the right MH (0.06 Hz, 95% CI [0.007, 0.12]). We further found an increased inhibition from the pregenual anterior cingulate cortex onto the bilateral MH (right hemisphere: 0.14 Hz, 95% CI [0.08, 0.20]; left hemisphere: 0.07 Hz, 95% CI [0.004, 0.14]) and a decreased inhibition from the middle cingulate onto the right MH (−0.01 Hz, 95% CI [-0.03, -0.01]). The left LH in turn received less inhibition from the angular gyrus (−0.01 Hz, 95% CI [-0.02, 0.01]) and exerted stronger inhibition on the substantia nigra (0.01 Hz, 95% CI [-0.007, 0.02]).

Outside the MH and LH, the substantia nigra pars compacta received a large number of inhibitory inputs from the areas of the cingulate, frontal and temporal cortices. Specifically, during the fasted as opposed to sated state, the substantia nigra pars compacta received a greater inhibition from the bilateral inferior temporal gyrus (left: 0.01 Hz, 95% CI [-0.007, 0.02]; right: 0.01 Hz, 95% CI [-0.007, 0.024]), and the pregenual anterior cingulate cortex (0.01 Hz, 95% CI [-0.02, 0.01]) as well as lower inhibition from the middle cingulate cortex (−0.02 Hz, 95% CI [-0.02, 0.006]) and inferior opercular frontal gyrus (−0.01 Hz, 95% CI [-0.02, 0.01]) (results are summarised in Table S1 and illustrated in Figure 4).

**Figure 4.**
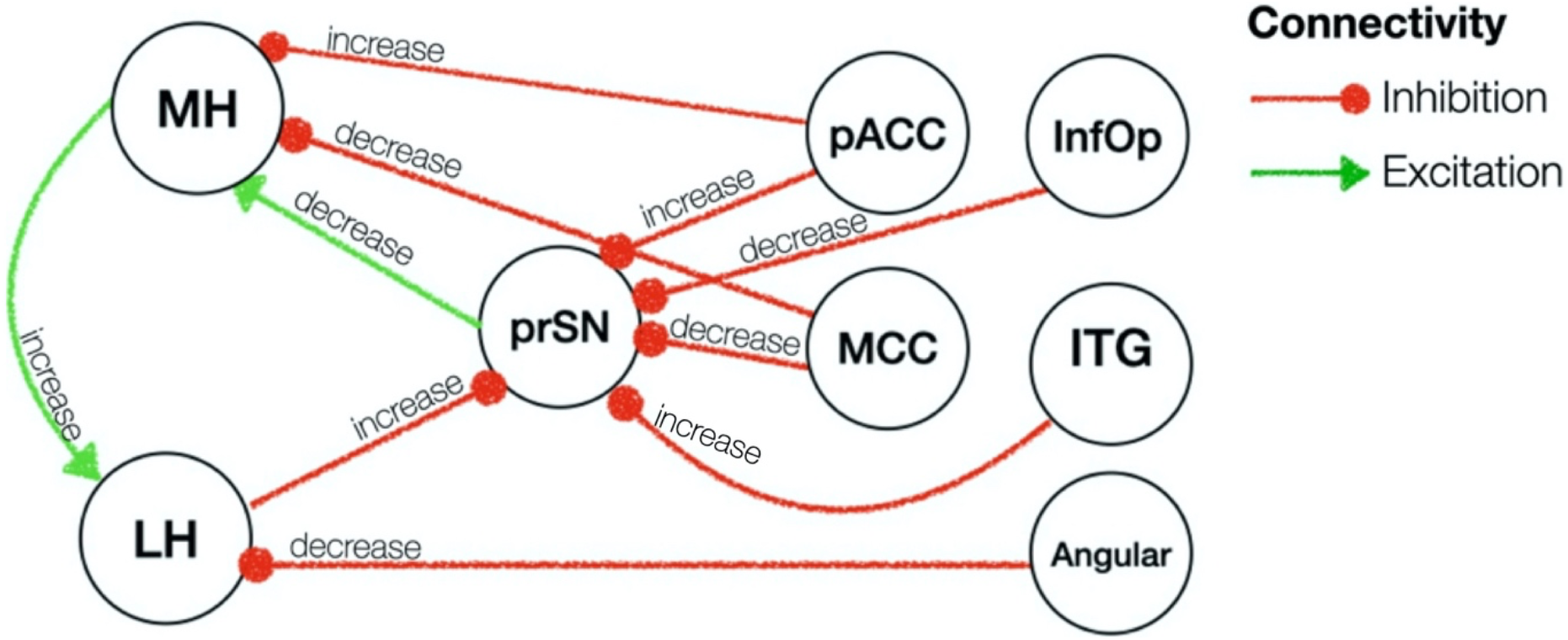
Effective connectivity of fasted vs. sated state. Red / green arrows indicate inhibitory / excitatory connectivity. All these connections had posterior probability of > 0.99 (very strong evidence). Abbreviations: prSN, substantia nigra pars compacta; MH, medial hypothalamus; LH, lateral hypothalamus; pACC, anterior cingulate cortex pregenual; MCC, middle cingulate and paracingulate gyri; ITG, inferior temporal gyrus; InfOp, inferior frontal gyrus, opercular part; Angular, Angular gyrus.

**Figure 5.**
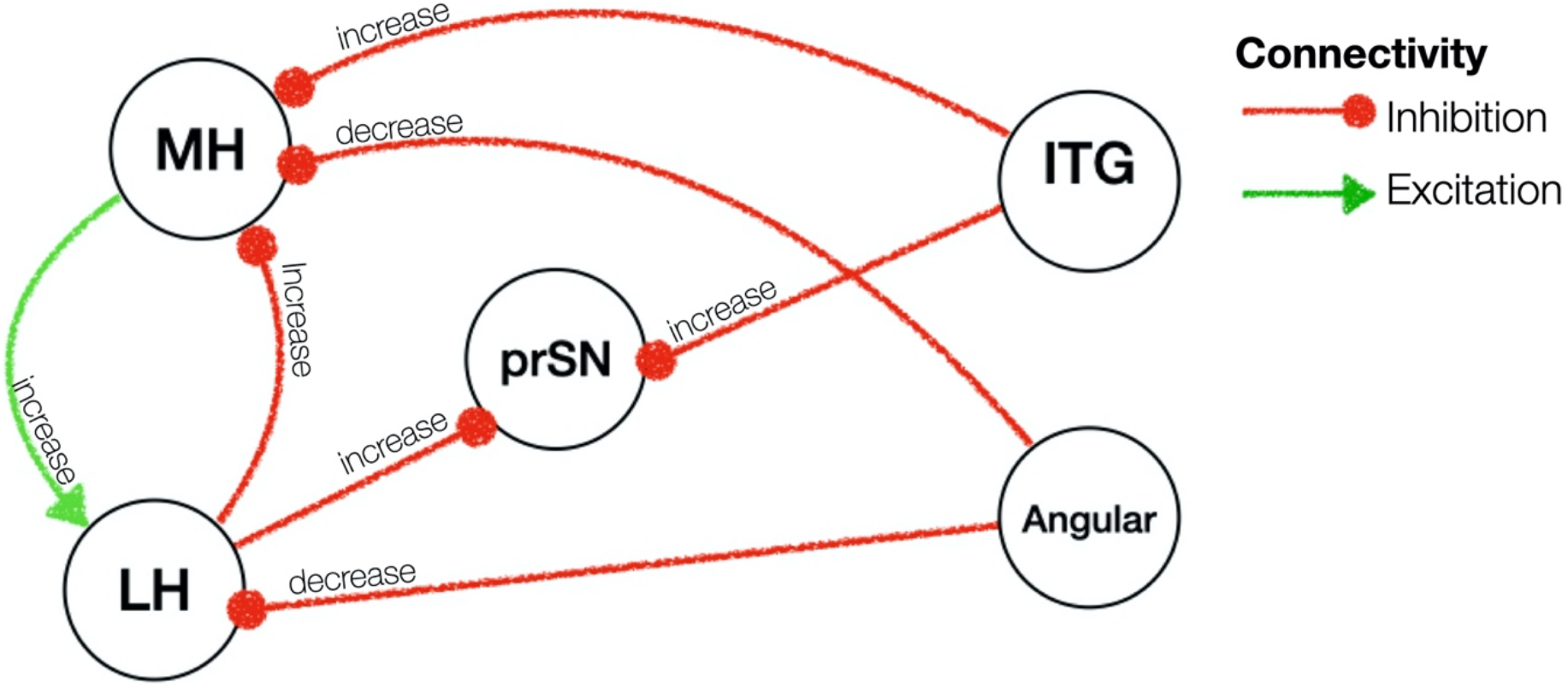
Effective connectivity of BMI. Red / green arrows indicate inhibitory / excitatory connectivity. Abbreviations: prSN, substantia nigra pars compacta; MH, medial hypothalamus; LH, lateral hypothalamus; ITG, inferior temporal gyrus; Angular, Angular gyrus.

### Effective connectivity changes of the MH and LH as a function of BMI

BMI, independent of homeostatic state, was associated with a greater excitatory influence from the left MH to the left LH (0.017 Hz, 95% CI [0.01, 0.03]) and greater inhibition from the right LH to the right MH (0.02 Hz, 95% CI [0.01, 0.02]). We further found a greater inhibition from the inferior temporal gyrus and angular gyrus to the MH (see Table 4 for lateralities and effect sizes). A greater inhibitory influence of the right LH on the substantia nigra (0.007 Hz, 95% CI [0.003, 0.012]) and the right MH (0.02 Hz, 95% CI [0.02, 0.024]) was also evident in individuals with a higher BMI. The substantia nigra pars compacta in turn received a greater inhibition from the bilateral inferior temporal gyrus (left: 0.014 Hz, 95% CI [0.01, 0.02]; right: 0.008 Hz, 95% CI [0.004, 0.01]) (results are summarised in Table S2 and illustrated in Figure 65).

### Effective connectivity changes in Fasted vs. Sated states in interaction with BMI

In the final analysis, we investigated how fasted-related connectivity changes were modulated by differences in BMI (Table 5 and Figure 6). During fasting relative to satiety, higher BMI was associated with a higher excitatory influence from the substantia nigra to the left MH (0.006 Hz, 95% CI [0.002, 0.009]). The left medial hypothalamus received an increased excitation from the right MH (0.013 Hz, 95% CI [0.004, 0.02]) as well as a decreased excitation from the left LH (−0.014 Hz, 95% CI [-0.02, -0.005]). The left MH also received higher inhibition from the anterior (0.001 Hz, 95% CI [0, 0.001]) and middle cingulate cortex (0.001 Hz, 95% CI [0, 0.002]). The substantia nigra pars compacta received a greater inhibition from the inferior opercular frontal gyrus (0.006 Hz, 95% CI [-0.01, -0.002]) (results are summarised in Table S3 and illustrated in Figure 6).

**Figure 6.**
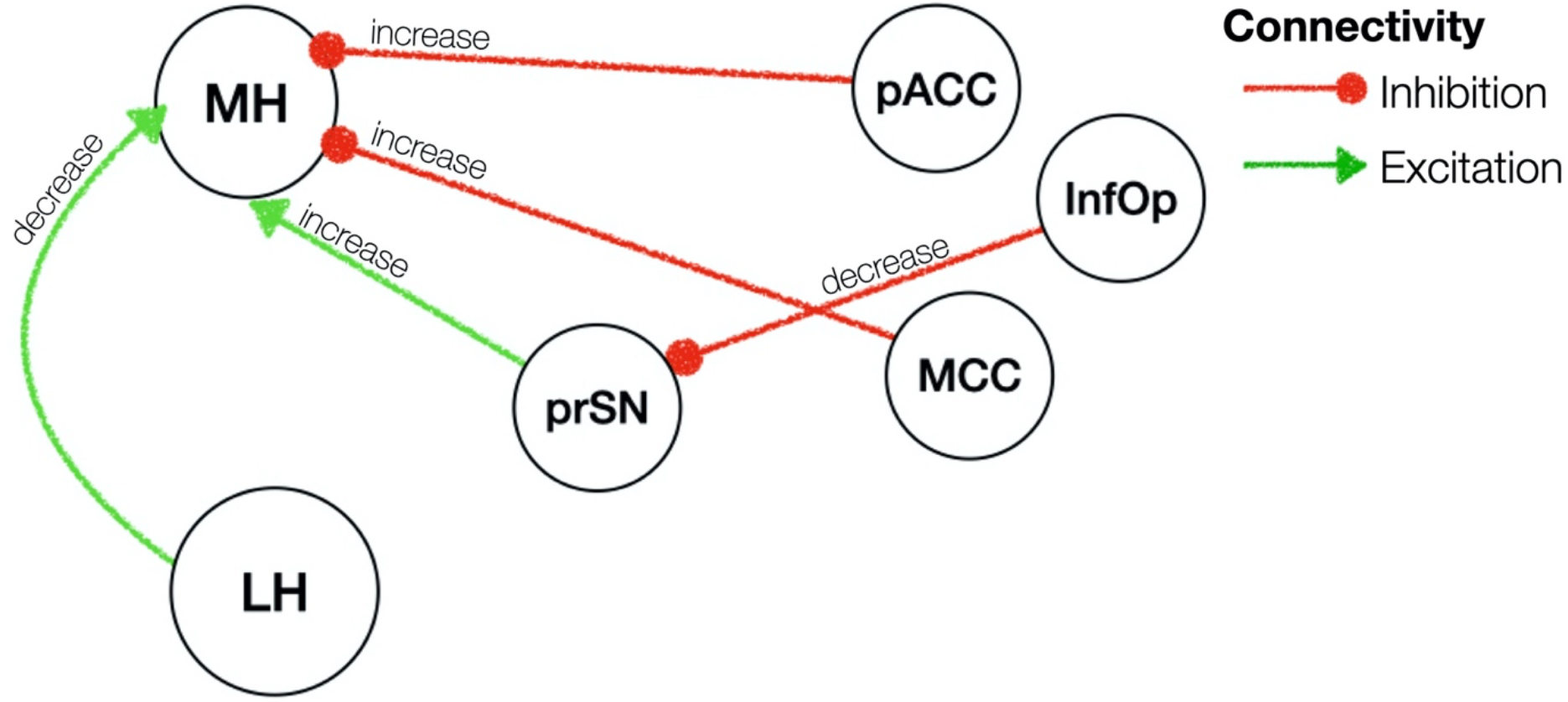
Effective Connectivity of BMI x Homeostatic state interaction effect. Red / green arrows indicate inhibitory / excitatory connectivity. Abbreviations: prSN, substantia nigra pars compacta; MH, medial hypothalamus; LH, lateral hypothalamus; pACC, anterior cingulate cortex pregenual; MCC, middle cingulate and paracingulate gyri; ITG, inferior temporal gyrus; InfOp, inferior frontal gyrus, opercular part.

## Discussion

This is the first study to reveal how the LH and MH causally interact with other neural regions and how their dynamics change with weight and energy state in humans. Adopting state of the art spectral dynamic causal modelling of resting-state fMRI data (Friston et al., 2014; Park et al., 2018; Razi et al., 2015), our results show two core networks interacting: (1) subcortical bidirectional connections between the LH, MH and the prSN, and (2) cortical top-down inhibition from frontal, cingulate and temporal onto the subcortical network. The prSN seems to represent a central hub interconnecting the subcortical and cortical neural systems. During the fasted compared to the sated state, regardless of weight status, we found increased inhibition between the LH and prSN as well as decreased inhibition between the prSN and MH, whereas the prSN received top-down inhibition from across the cortex, which may represent an adaptive motivational drive to seek food while hungry, in fitting with animal studies (Cassidy & Tong, 2017; Nieh et al., 2016; Rossi et al., 2019; Rossi & Stuber, 2018). However, individuals with excess weight revealed a similar hypothalamic network communication irrespective of being in a fasted or sated state. Further, when taking into consideration excess weight, they showed a reverse communication pattern of decreased substantia nigra-MH inhibition during the fasted state. The neural network communications involved in the regular processes of food seeking after fasting may therefore be disrupted in individuals with excess weight, providing a compelling hypothesis for food overconsumption beyond metabolic needs.

Our results from whole-brain functional connectivity analyses largely reflect those reported in earlier neuroimaging studies (Kullmann et al., 2014; Martín-Pérez et al., 2019; Zhang et al., 2018). Both resting-state activity of LH and the MH was correlated with resting-state activity of prSN, middle and anterior cingulate cortex, inferior frontal and temporal gyrus as well as angular gyrus. Consistent with animal and human research, the LH was more strongly connected than the MH across the entire brain ranging from subcortical areas (e.g., red nucleus, putamen) to all neocortical areas. This finding strengthens the established role of the LH as a central interface integrating diverse central and peripheral signals through a complex large-scale neural network that may coordinate adaptive behavioural responses related to motivation and controlled feeding behaviour (Bonnavion et al., 2016; Petrovich, 2018). The MH in turn was more strongly connected to the prSN, cerebellum and precuneus. These and previous findings (Kullmann et al., 2014; Martín-Pérez et al., 2019; Zhang et al., 2018) highlight the potential of a dual hypothalamic functionality resulting from distinct LH and MH neural networks. However, these characterisations have been limited as the directionality and valence of the interactions within these networks have remained unknown. Here, we have extended the characterisation of these network by means of spDCM investigating the directed communication of the LH and MH network and their changes as a function of both BMI and energy state.

Irrespective of BMI and energy state, the prSN emerges as a key area connecting the hypothalamus with neocortical regions. The prSN processes autonomic, gut-induced rewards regulating motivational and emotional states (e.g., Gutierrez et al., 2020; Han et al., 2018). Hormones implicated in regulating the homeostatic system also impinge directly on dopamine neurons in the prSN (Palmiter, 2007). Anatomically and functionally the prSN is highly interconnected with the ventral tegmental area (VTA) and both regions contribute to motivation and reward processing (Ilango et al., 2014; Kwon & Jang, 2014). In hungry mice, inhibitory inputs from the LH to the VTA inhibited dopamine release, resulting in increased motivation to seek and approach food (Nieh et al., 2016). In this study we found an increased inhibitory influence from the LH to the prSN when participants were hungry. Given the strong prSN-VTA interconnectivity and interchangeable functionality (Ilango et al., 2014; Kwon & Jang, 2014), it is reasonable to assume that the increased inhibition from the LH to the prSN in humans might mirror a necessary survival mechanism to increase appetitive motivation to prevent starvation. Notably, in individuals with higher BMI, regardless of their energy state, this inhibition from LH to prSN persisted. This failure to ‘shut-off’ the inhibitory signalling might reflect an underlying neural trigger for increased motivation for food regardless of homeostatic state (Berthoud, 2004, 2012; Berthoud et al., 2017; Cassidy & Tong, 2017).

During the fasted compared to the sated state, we also found a decreased excitation from the prSN to the MH. The MH contains a diverse array of nuclei and circuits, including the anorectic melanocortin system that reduces food consumption, as well as increasing energy expenditure (Kühnen et al., 2019). Thus, the reduced excitation during fasting compared to satiety may reflect reduced activation of this anorectic pathway. However, it is also important to note that the MH contains strong drivers of appetite and motivation. Agouti-related peptide neurons in the hypothalamic arcuate nucleus drive food intake and motivation (Andermann & Lowell, 2017). In individuals with excess weight, this excitatory connectivity between prSN and MH increased during the fasted compared to the sated state. This might contribute to increased food seeking and consumption in response to energy deprivation among individuals with overweight/obesity (Kühnen et al., 2019). Clearly, future research in humans is required to examine the activity of specific hypothalamic nuclei within the MH region in relation to food seeking and obesity; however, this is currently beyond the technical capability of MRI.

In addition to the communication between MH and LH with the prSN, we also found intra-hypothalamic connectivity between the LH and MH across weight and homeostatic state. The dynamic from the ventral MH to the LH has been previously observed in animals (Canteras et al., 1994; Horst & Luiten, 1986; Luiten et al., 1980); however, its functional significance remains untested and further studies in humans are needed to elucidate the functionality of intra-hypothalamic connectivity patterns.

During the fasted as opposed to the sated state, we found an increased top-down inhibition from the pregenual anterior cingulate cortex to the MH, yet a decreased top-down inhibition from the middle cingulate gyrus to the MH and prSN. While it is not possible to definitively disentangle the role of these network dynamics in the current context, previous neuroimaging studies that have dissociated the functions of the pregenual anterior cingulate cortex and the middle cingulate gyrus provide for some conjecture (Stevens, 2011). In particular, spontaneous activity in the pregenual anterior cingulate cortex is associated with affective processing, and anticorrelated with activity in sensorimotor areas. In contrast, activity in the middle cingulate gyrus is temporally coupled with activity in sensorimotor areas, and functionally connected with areas involved in cognitive control. The causal dynamics we report herein between the middle cingulate gyrus, pregenual anterior cingulate cortex and MH might therefore suggest interactions between homeostatic inputs and affective, sensorimotor and cognitive networks dynamics. Note that the increased inhibition from the pregenual anterior cingulate cortex to the MH was not related to BMI. However, increased inhibition from the middle cingulate gyrus to the MH was exacerbated in participants with higher BMIs. Future studies are needed to clarify if this alteration is associated with core symptoms of obesity.

In addition to the cingulate cortex, the angular gyrus had a decreased impact on the LH, when individuals were fasted or in individuals with excess weight. Furthermore, all cortical areas inhibited either the prSN or hypothalamus irrespective of weight and homeostatic state. Whereas the cingulate cortex is a hub for sensory, motivational and cognitive information, the prefrontal and parietal cortex are more predominantly associated with executive control (Seeley et al., 2007). In participants with excess weight, a differential pattern within the executive control network has been observed in fMRI activation studies using food stimuli (Franssen et al., 2020). Recently, it has also been shown that obesity is related to prominent functional connectivity alterations mainly in prefrontal regions during resting-state as well as in response to food stimuli (García-García et al., 2013; Kullmann et al., 2012). Thus, our resting-state findings might further add to the possibility of disrupted communication between the executive control network and regions regulating metabolic needs in individuals with excess weight.

In the light of the proposed mechanisms here, we note, however, that the relationship between resting-state effective connectivity and its cognitive correlates remains elusive. The inter-individual variations in effective connectivity do not necessarily overlap with the inter-individual variations in effective connectivity during task performance (Fox & Raichle, 2007; Jung et al., 2018). At this stage, only one study has revealed that resting-state effective connectivity might facilitate task performance but may not reflect task-based network dynamics (Jung et al., 2018).Future studies are needed to address whether the resting-state dynamics revealed in our study are also engaged during task performance and how potential deviations might translate to differences in behaviour or clinical phenotypes.

In conclusion our study provides new insights into how hunger and satiety states affect hypothalamic circuit dynamics, which involve a distributed network of midbrain and cortical areas with a key role of the substantia nigra pars compacta. We also identified unique aspects of network organisation associated with obesity, which involve the reciprocal connections between the lateral and medial hypothalamus, and the input from the substantia nigra to the medial hypothalamus.

## Acknowledgements

The authors thank Richard McIntyre, Naomi Kakoschke, Amelia Romei, Tori Gaunson and Tiffany Falcone for help with MRI data acquisition. This study was supported by a NHMRC grant (1140197) granted to Antonio Verdejo-Garcia, Zane Andrews, and Ian Harding.

## Data Availability Statement

Data for main spectral dynamic causal modelling are available on github: https://github.com/KatharinaVoigt1/spDCM_Hypo.git.

### Appendix

#### Supplementary Analyses

##### Relationship between BMI and Subjective Hunger Reports

There was no significant relationship between BMI and reported subjective hunger during the fasted (M = 4.42, SD = 1.44) and sated condition (M = 3.22, SD = 1.48) (Main effect of subjective hunger on BMI: *β* = -0.06, *SE* = 0.18, *p* = .76; Interaction effect between subjective hunger and homeostatic condition on BMI: *β* = 0.17, *SE* = 0.25, *p* = .51) (Figure S1). When excluded an outlier based on BMI (ID_1 = 55.56 kg/m2), this relationship remained.

**Figure S1.**
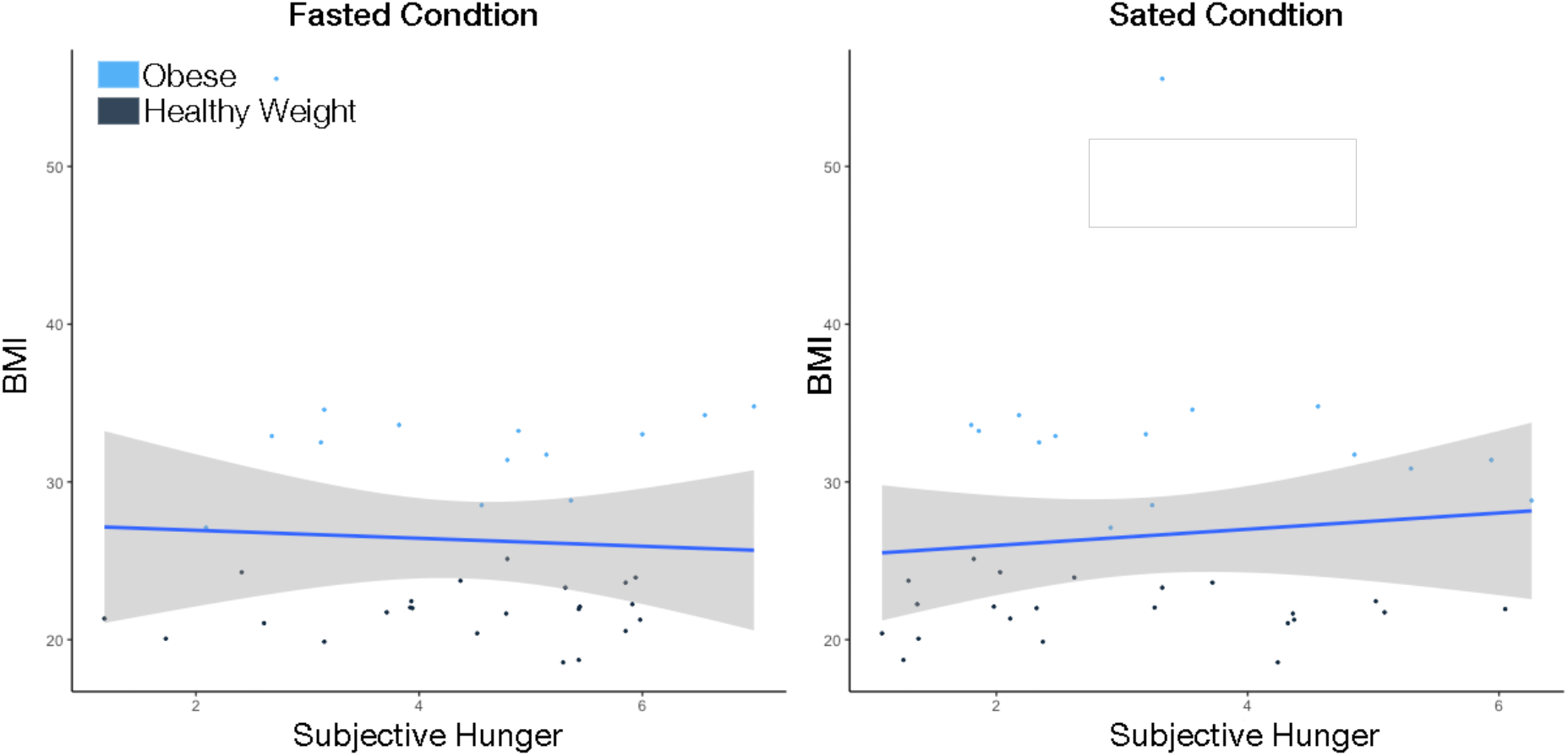
Relationship between reported hunger and BMI across the fasted and sated condition. Grey area indicates 95% confidence interval of linear model’s predictions.

##### Relationship between BMI and Fasting Blood Glucose Levels

There was a significant positive relationship between BMI and fasting blood glucose levels: with each increase in BMI by 1 kg/m^2^, blood glucose levels increased by 0.31 mg/dL (*β* = 0.31, *SE* = 0.15, *p* = .04) (Figure S2, left panel). However, when excluding outliers in BMI (ID_1 = 55.56 kg/m2) and blood glucose levels (ID_18 = 6.4 mg/dL; ID_23 = 8.4 mg/dL), this relationship did not persist (Figure S2, right panel), (*β* = 0.27, *SE* = 0.28, *p* = .33). In order to avoid potentially extracting meaningful network information by excluding these participants, we instead controlled for fasting blood glucose levels when investigating the dynamics of the hypothalamic neural network model in relation with BMI (i.e. Model 2, Model 3, and Model 4).

**Figure S2.**
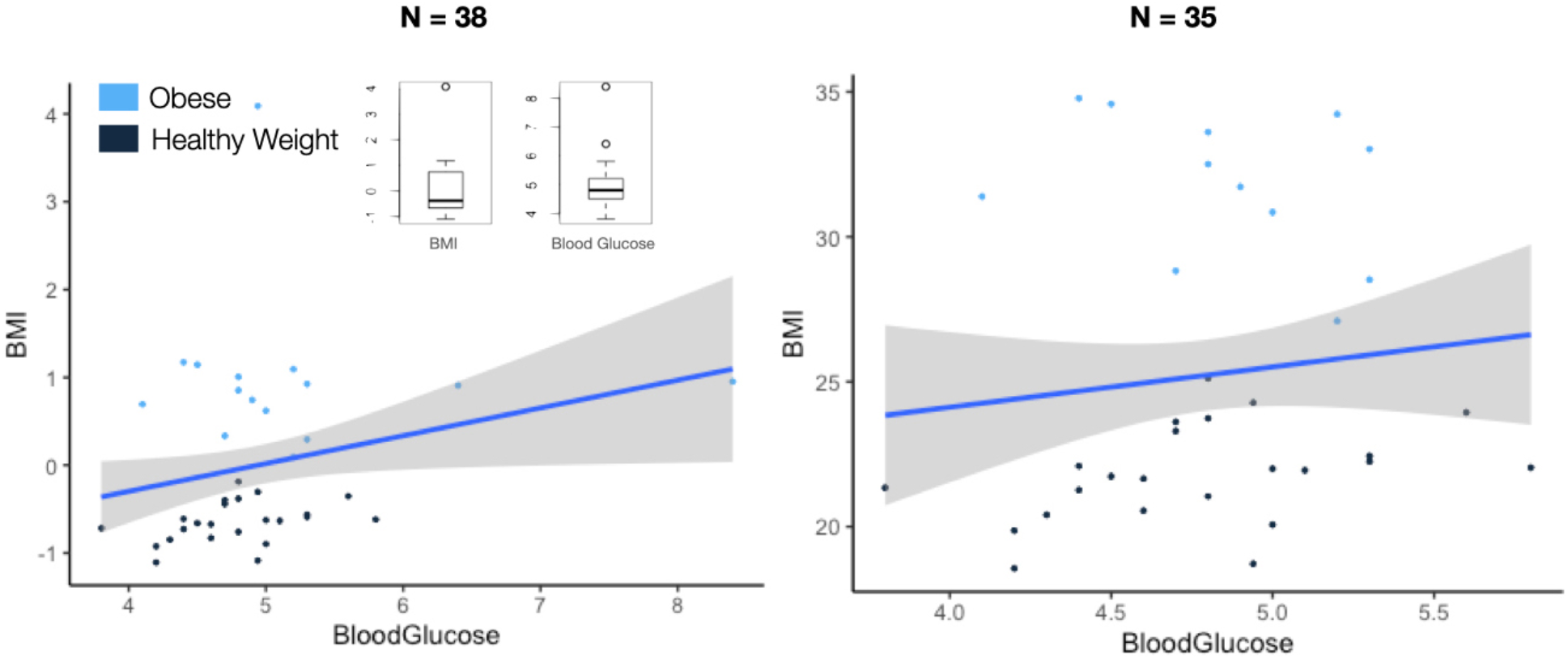
Relationship between BMI and fasting blood glucose levels for all participants (N = 38, left panel) and for when outliers were excluded (N = 35, right panel). Grey area indicates 95% confidence interval of linear model’s predictions.

#### Supplementary Results

**Table S1.**
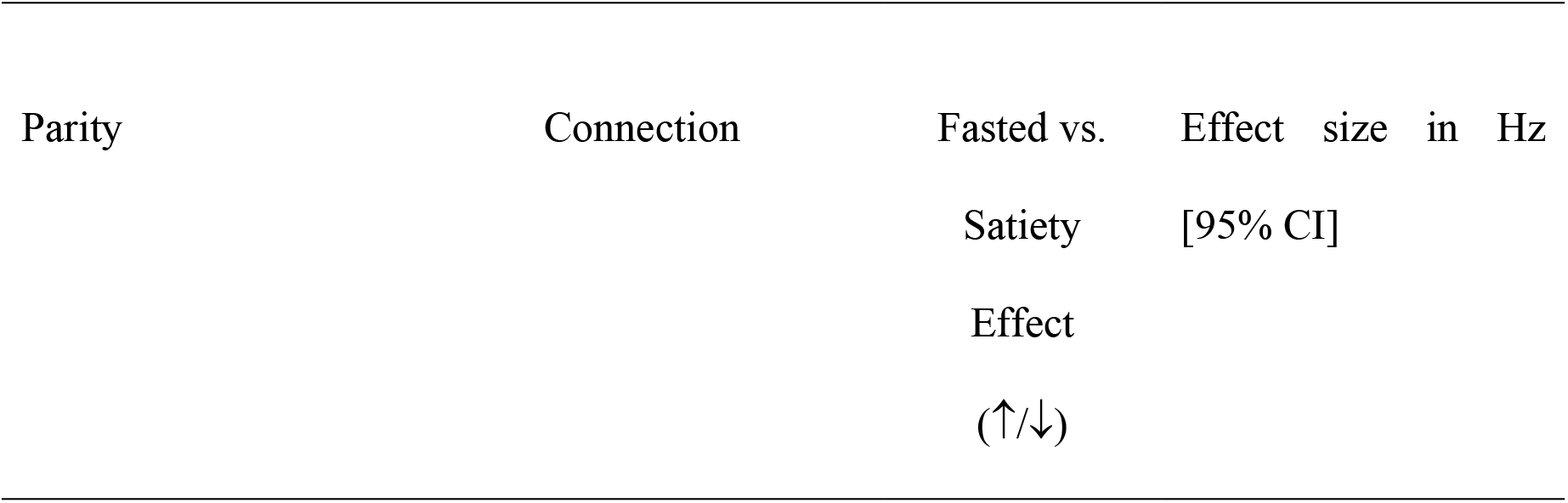
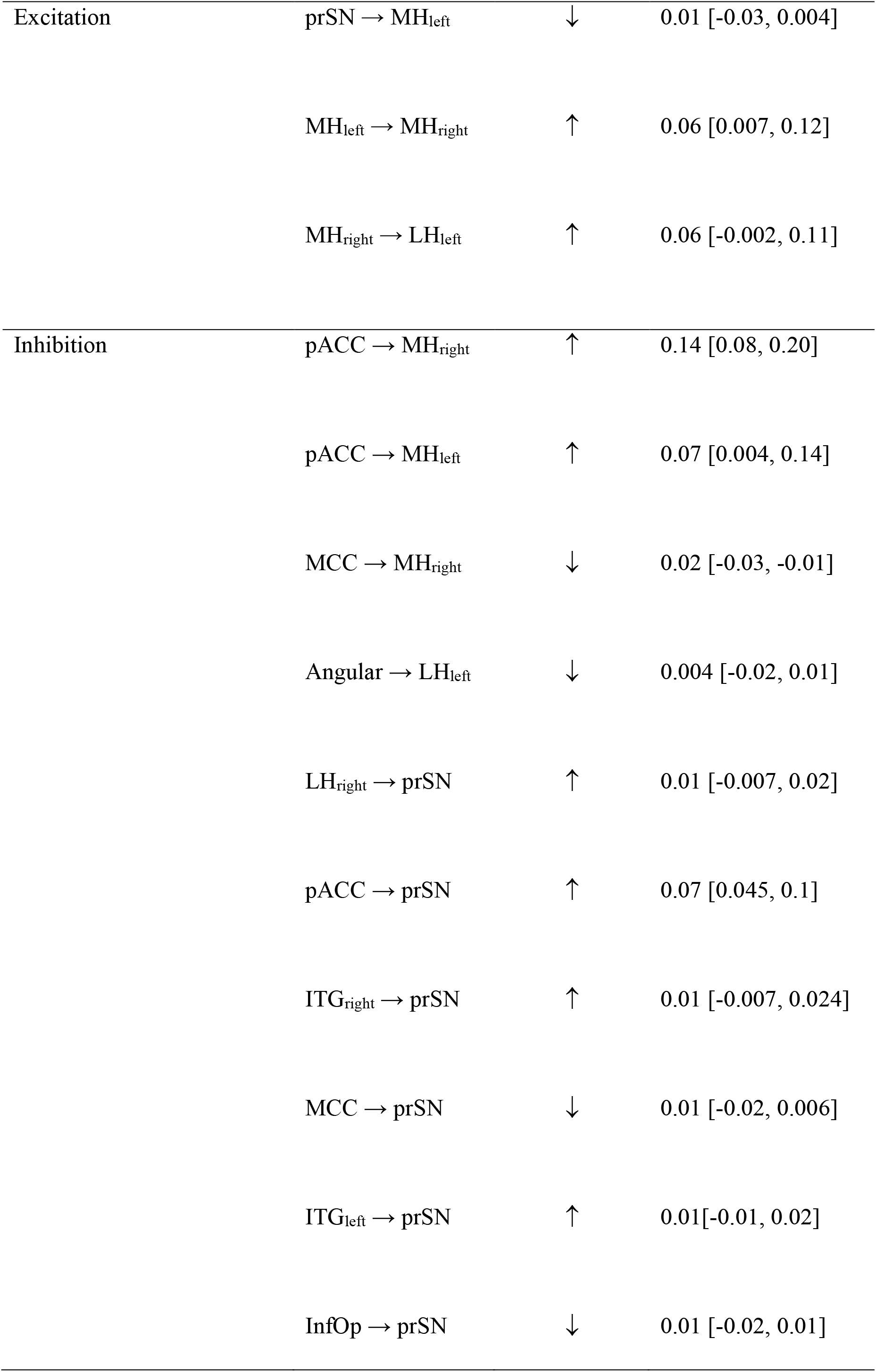
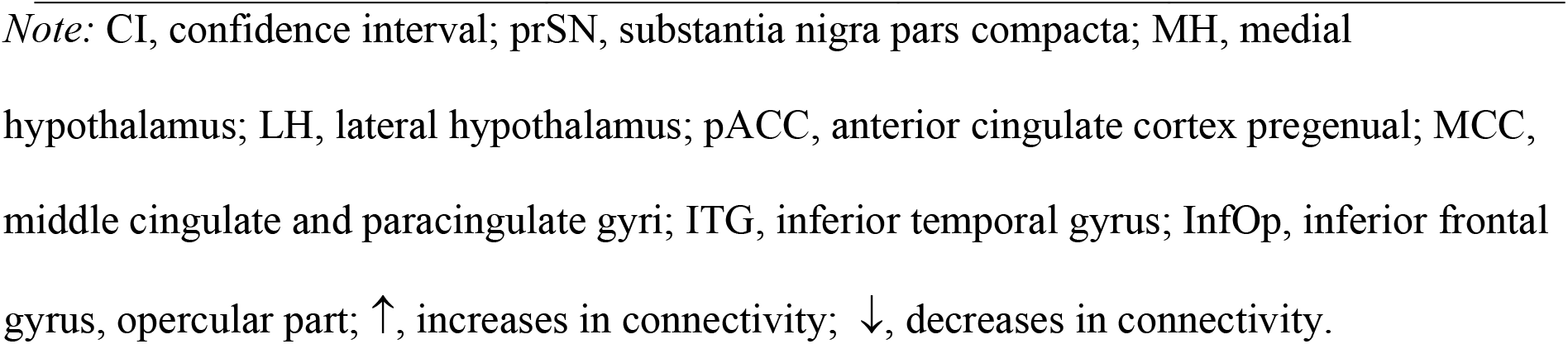
Modulation of the Hypothalamic Network by Homeostatic State (Fasted vs. Sated)

|  |  |  |  |
| --- | --- | --- | --- |
| Excitation | $\text{prSN} \rightarrow \text{MH}_{\text{left}}$ | ↓ | 0.01 [-0.03, 0.004] |
| | $\text{MH}_{\text{left}} \rightarrow \text{MH}_{\text{right}}$ | ↑ | 0.06 [0.007, 0.12] |
| | $\text{MH}_{\text{right}} \rightarrow \text{LH}_{\text{left}}$ | ↑ | 0.06 [-0.002, 0.11] |
| Inhibition | $\text{pACC} \rightarrow \text{MH}_{\text{right}}$ | ↑ | 0.14 [0.08, 0.20] |
| | $\text{pACC} \rightarrow \text{MH}_{\text{left}}$ | ↑ | 0.07 [0.004, 0.14] |
| | $\text{MCC} \rightarrow \text{MH}_{\text{right}}$ | ↓ | 0.02 [-0.03, -0.01] |
| | $\text{Angular} \rightarrow \text{LH}_{\text{left}}$ | ↓ | 0.004 [-0.02, 0.01] |
| | $\text{LH}_{\text{right}} \rightarrow \text{prSN}$ | ↑ | 0.01 [-0.007, 0.02] |
| | $\text{pACC} \rightarrow \text{prSN}$ | ↑ | 0.07 [0.045, 0.1] |
| | $\text{ITG}_{\text{right}} \rightarrow \text{prSN}$ | ↑ | 0.01 [-0.007, 0.024] |
| | $\text{MCC} \rightarrow \text{prSN}$ | ↓ | 0.01 [-0.02, 0.006] |
| | $\text{ITG}_{\text{left}} \rightarrow \text{prSN}$ | ↑ | 0.01 [-0.01, 0.02] |
| | $\text{InfOp} \rightarrow \text{prSN}$ | ↓ | 0.01 [-0.02, 0.01] |

*Note:* CI, confidence interval; prSN, substantia nigra pars compacta; MH, medial hypothalamus; LH, lateral hypothalamus; pACC, anterior cingulate cortex pregenual; MCC, middle cingulate and paracingulate gyri; ITG, inferior temporal gyrus; InfOp, inferior frontal gyrus, opercular part; ↑, increases in connectivity; ↓, decreases in connectivity.

**Table S2.**
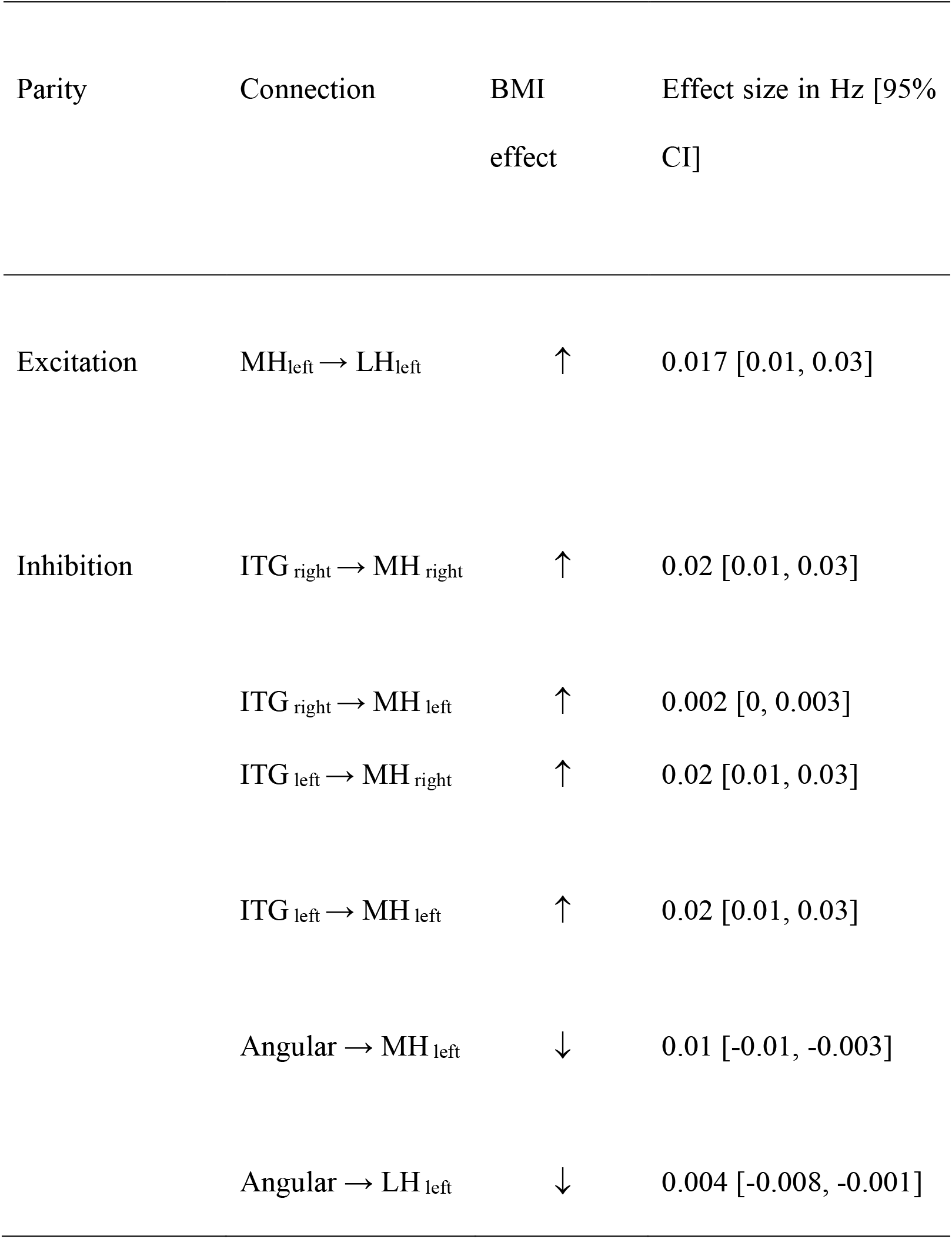
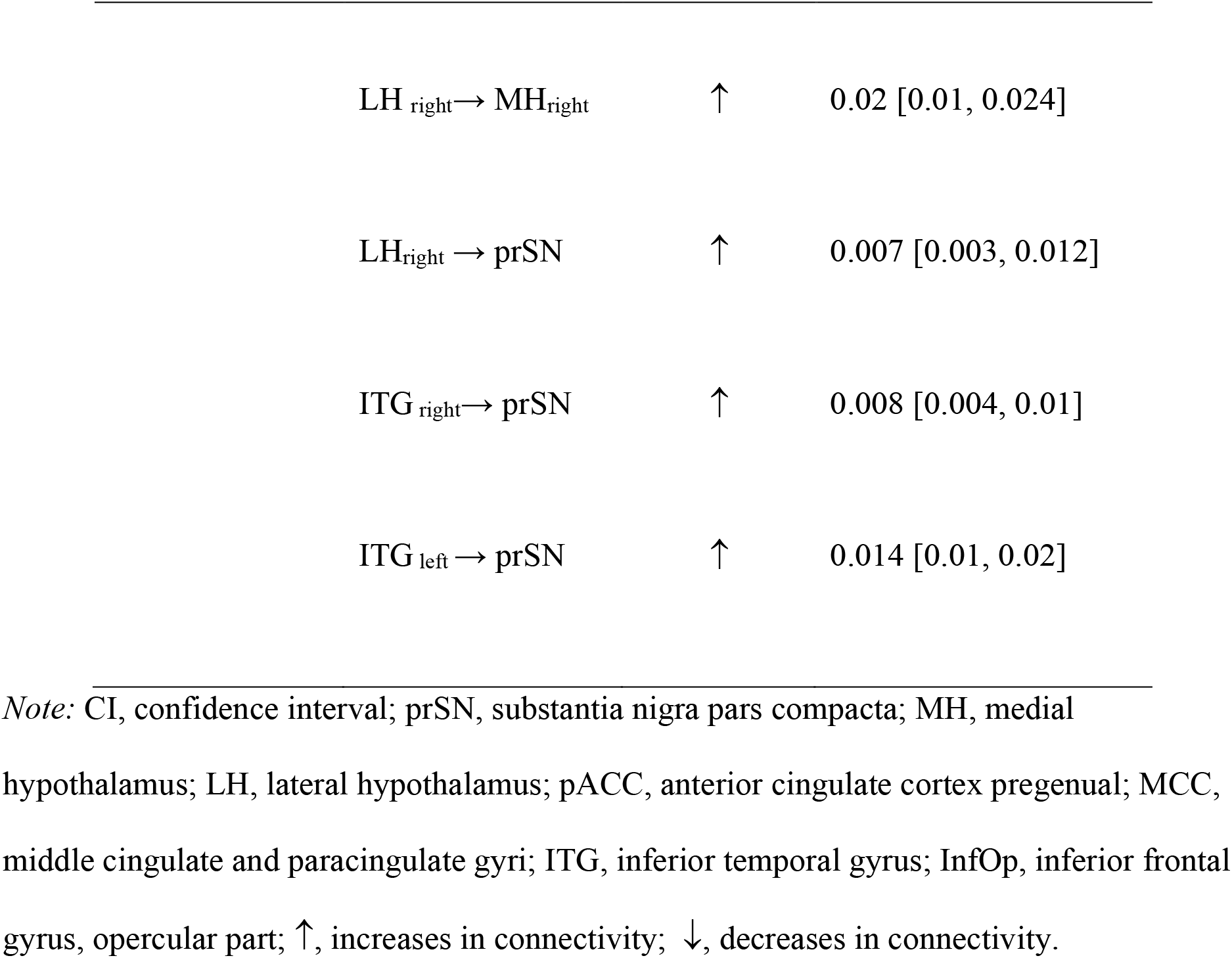
Differences in the Hypothalamic Network associated with BMI

| Parity | Connection | BMI effect | Effect size in Hz [95% CI] |
| --- | --- | --- | --- |
| Excitation | MH<sub>left</sub> → LH<sub>left</sub> | ↑ | 0.017 [0.01, 0.03] |
| Inhibition | ITG<sub>right</sub> → MH<sub>right</sub> | ↑ | 0.02 [0.01, 0.03] |
|  | ITG<sub>right</sub> → MH<sub>left</sub> | ↑ | 0.002 [0, 0.003] |
|  | ITG<sub>left</sub> → MH<sub>right</sub> | ↑ | 0.02 [0.01, 0.03] |
|  | ITG<sub>left</sub> → MH<sub>left</sub> | ↑ | 0.02 [0.01, 0.03] |
|  | Angular → MH<sub>left</sub> | ↓ | 0.01 [-0.01, -0.003] |
|  | Angular → LH<sub>left</sub> | ↓ | 0.004 [-0.008, -0.001] |

|  |  |  |
| --- | --- | --- |
| LH<sub>right</sub> → MH<sub>right</sub> | ↑ | 0.02 [0.01, 0.024] |
| LH<sub>right</sub> → prSN | ↑ | 0.007 [0.003, 0.012] |
| ITG<sub>right</sub> → prSN | ↑ | 0.008 [0.004, 0.01] |
| ITG<sub>left</sub> → prSN | ↑ | 0.014 [0.01, 0.02] |

*Note:* CI, confidence interval; prSN, substantia nigra pars compacta; MH, medial hypothalamus; LH, lateral hypothalamus; pACC, anterior cingulate cortex pregenual; MCC, middle cingulate and paracingulate gyri; ITG, inferior temporal gyrus; InfOp, inferior frontal gyrus, opercular part; ↑, increases in connectivity; ↓, decreases in connectivity.

**Table S3.**
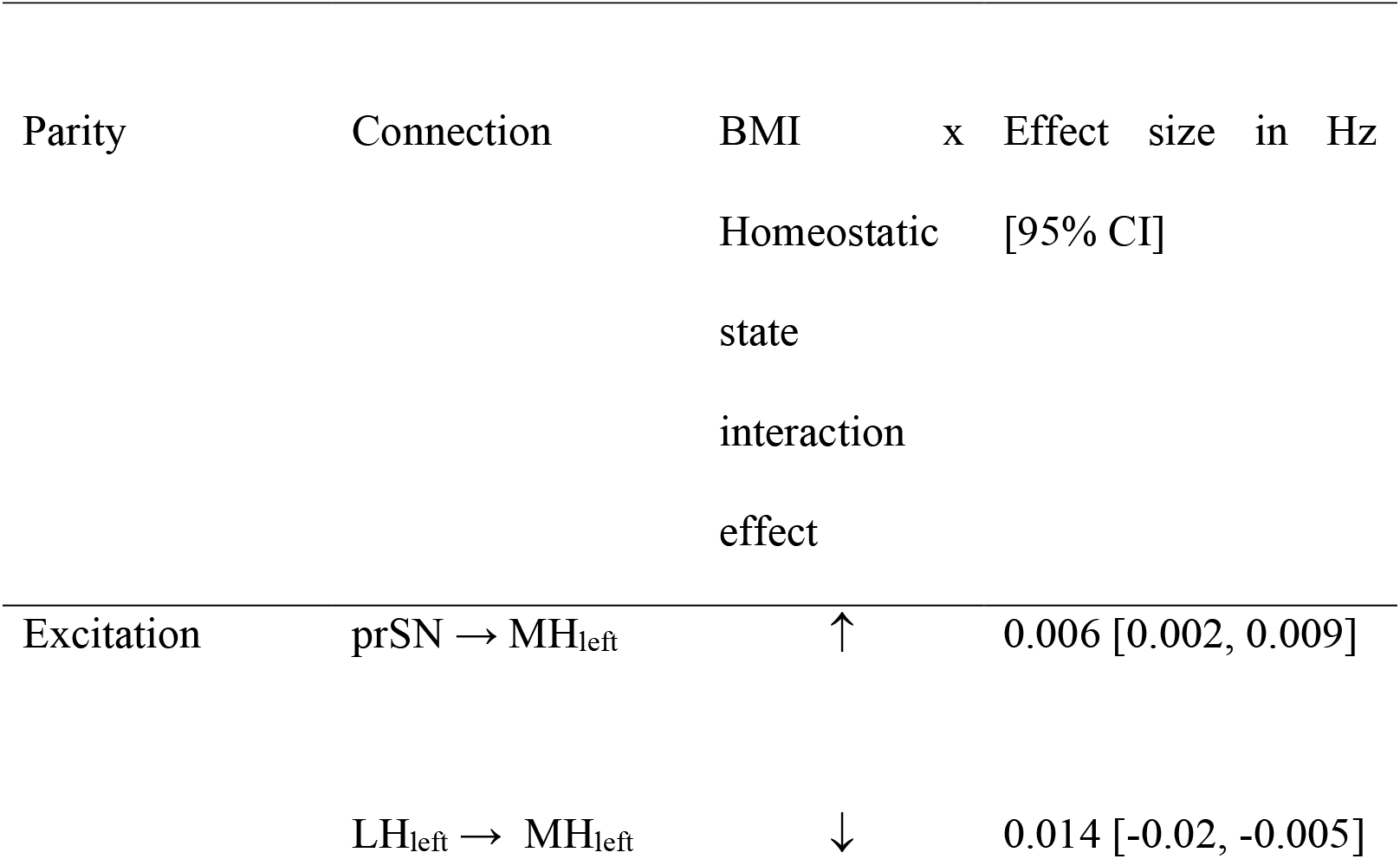
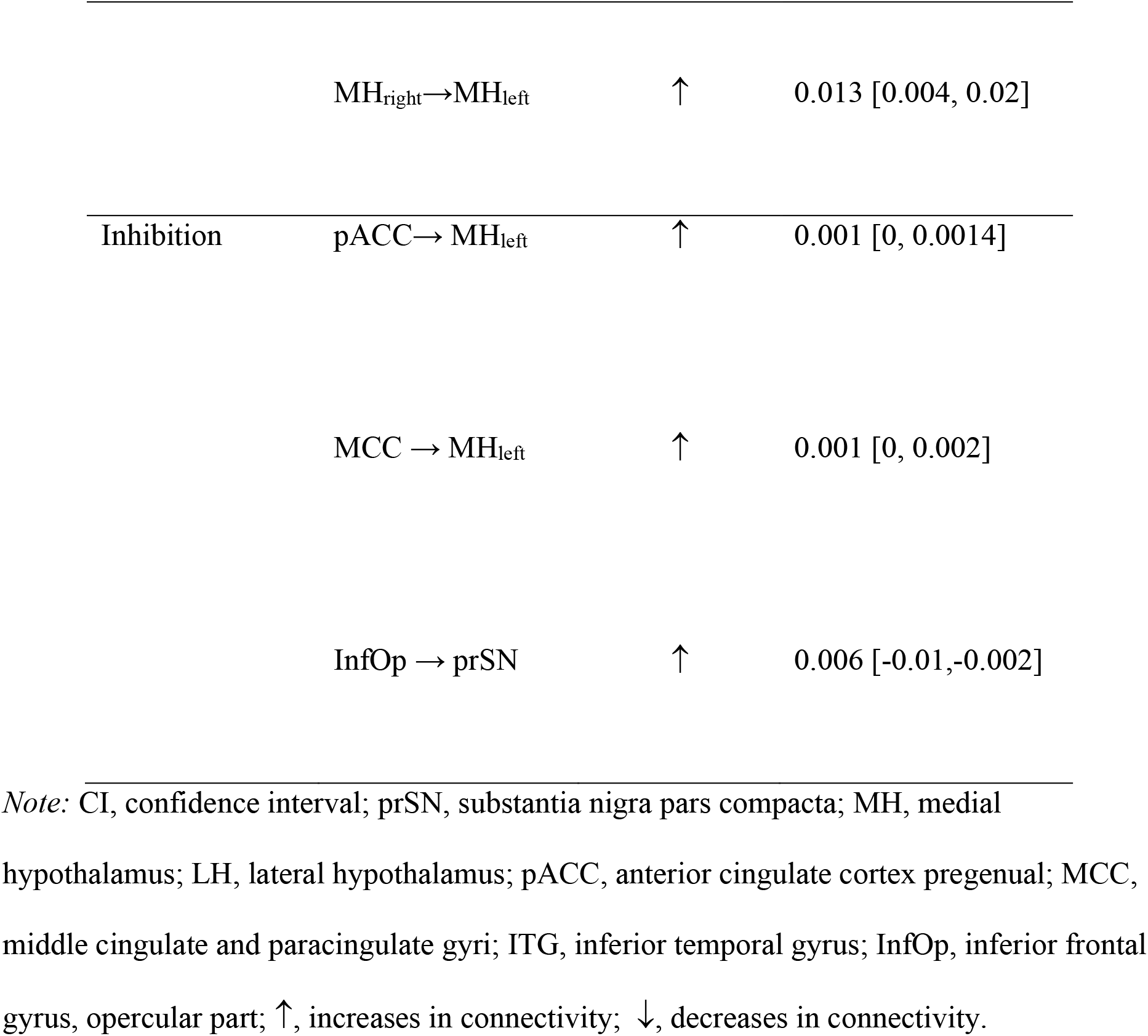
Hypothalamic network associated with BMI x Homeostatic State interaction

| Parity | Connection | BMI x Homeostatic state interaction effect | Effect size in Hz [95% CI] |
| --- | --- | --- | --- |
| Excitation | prSN → MH<sub>left</sub> | ↑ | 0.006 [0.002, 0.009] |
|  | LH<sub>left</sub> → MH<sub>left</sub> | ↓ | 0.014 [-0.02, -0.005] |
|  | MH<sub>right</sub>→MH<sub>left</sub> | ↑ | 0.013 [0.004, 0.02] |
| Inhibition | pACC→ MH<sub>left</sub> | ↑ | 0.001 [0, 0.0014] |
|  | MCC → MH<sub>left</sub> | ↑ | 0.001 [0, 0.002] |
|  | InfOp → prSN | ↑ | 0.006 [-0.01,-0.002] |

*Note:* CI, confidence interval; prSN, substantia nigra pars compacta; MH, medial hypothalamus; LH, lateral hypothalamus; pACC, anterior cingulate cortex pregenual; MCC, middle cingulate and paracingulate gyri; ITG, inferior temporal gyrus; InfOp, inferior frontal gyrus, opercular part; ↑, increases in connectivity; ↓, decreases in connectivity.

## References

Anand, B. K., & Brobeck, J. R. (1951). Hypothalamic control of food intake in rats and cats. The Yale Journal of Biology and Medicine, 24(2), 123–140. PubMed.

Andermann, M. L., & Lowell, B. B. (2017). Toward a Wiring Diagram Understanding of Appetite Control. Neuron, 95(4), 757–778. https://doi.org/10.1016/j.neuron.2017.06.014

Baroncini, M., Jissendi, P., Balland, E., Besson, P., Pruvo, J.-P., Francke, J.-P., Dewailly, D., Blond, S., & Prevot, V. (2012). MRI atlas of the human hypothalamus. NeuroImage (Orlando, Fla.), 59(1), 168–180. https://doi.org/10.1016/j.neuroimage.2011.07.013

Berthoud, H.-R. (2004). Neural control of appetite: Cross-talk between homeostatic and non-homeostatic systems. Appetite, 43(3), 315–317. https://doi.org/10.1016/j.appet.2004.04.009

Berthoud, H.-R. (2012). The neurobiology of food intake in an obesogenic environment. Proceedings of the Nutrition Society, 71(4), 478–487. https://doi.org/10.1017/S0029665112000602

Berthoud, H.-R., Münzberg, H., & Morrison, C. D. (2017). Blaming the Brain for Obesity: Integration of Hedonic and Homeostatic Mechanisms. Gastroenterology, 152(7), 1728–1738. https://doi.org/10.1053/j.gastro.2016.12.050

Bonnavion, P., Mickelsen, L. E., Fujita, A., de Lecea, L., & Jackson, A. C. (2016). Hubs and spokes of the lateral hypothalamus: Cell types, circuits and behaviour: LHA cell types and circuits. The Journal of Physiology, 594(22), 6443–6462. https://doi.org/10.1113/JP271946

Brobeck, J. R., Tepperman, J., & Long, C. N. (1943). Experimental Hypothalamic Hyperphagia in the Albino Rat. The Yale Journal of Biology and Medicine, 15(6), 831–853. PubMed.

Burdakov, D., & Karnani, M. M. (2020). Ultra-sparse Connectivity within the Lateral Hypothalamus. Current Biology, 30(20), 4063-4070.e2. https://doi.org/10.1016/j.cub.2020.07.061

Canteras, N. S., Simerly, R. B., & Swanson, L. W. (1994). Organization of projections from the ventromedial nucleus of the hypothalamus: A Phaseolus vulgaris-Leucoagglutinin study in the rat. Journal of Comparative Neurology, 348(1), 41–79. https://doi.org/10.1002/cne.903480103

Cassidy, R. M., & Tong, Q. (2017). Hunger and Satiety Gauge Reward Sensitivity. Frontiers in Endocrinology, 8. https://doi.org/10.3389/fendo.2017.00104

Chen, R., Wu, X., Jiang, L., & Zhang, Y. (2017). Single-Cell RNA-Seq Reveals Hypothalamic Cell Diversity. Cell Reports, 18(13), 3227–3241. https://doi.org/10.1016/j.celrep.2017.03.004

Elmquist, J. K., Elias, C. F., & Saper, C. B. (1999). From lesions to leptin: Hypothalamic control of food intake and body weight. Neuron, 22(2), 221.

Florent, V., Baroncini, M., Jissendi-Tchofo, P., Lopes, R., Vanhoutte, M., Rasika, S., Pruvo, J.-P., Vignau, J., Verdun, S., Johansen, J. E., Pigeyre, M., Bouret, S. G., Nilsson, I. A. K., & Prevot, V. (2020). Hypothalamic Structural and Functional Imbalances in Anorexia Nervosa. Neuroendocrinology, 110(6), 552. https://doi.org/10.1159/000503147

Fox, M. D., & Raichle, M. E. (2007). Spontaneous fluctuations in brain activity observed with functional magnetic resonance imaging. Nature Reviews Neuroscience, 8(9), 700–711. https://doi.org/10.1038/nrn2201

Franssen, S., Jansen, A., Schyns, G., van den Akker, K., & Roefs, A. (2020). Neural Correlates of Food Cue Exposure Intervention for Obesity: A Case-Series Approach. Frontiers in Behavioral Neuroscience, 14, 46. https://doi.org/10.3389/fnbeh.2020.00046

Friston, K., Harrison, L., & Penny, W. (2003). Dynamic causal modelling. NeuroImage, 19(4), 97–109.

Friston, K. J., Litvak, V., Oswal, A., Razi, A., Stephan, K. E., van Wijk, B. C. M., Ziegler, G., & Zeidman, P. (2016). Bayesian model reduction and empirical Bayes for group (DCM) studies. NeuroImage, 128, 413–431. https://doi.org/10.1016/j.neuroimage.2015.11.015

Friston, K., Kahan, J., Biswal, B., & Razi, A. (2014). A DCM for resting state fMRI. NeuroImage, 94, 396–407.

Friston, K., Mattout, J., Trujillo-Barreto, N., Ashburner, J., & Penny, W. (2007). Variational free energy and the Laplace approximation. NeuroImage, 34(1), 220–234. https://doi.org/10.1016/j.neuroimage.2006.08.035

Gabriela Pop, M., Crivii, C., & Opincariu, I. (2018). Anatomy and Function of the Hypothalamus. In S. J. Baloyannis & J. Oxholm Gordeladze (Eds.), Hypothalamus in Health and Diseases. IntechOpen. https://doi.org/10.5772/intechopen.80728

García-García, I., Jurado, M. Á., Garolera, M., Segura, B., Sala-Llonch, R., Marqués-Iturria, I., Pueyo, R., Sender-Palacios, M. J., Vernet-Vernet, M., Narberhaus, A., Ariza, M., & Junqué, C. (2013). Alterations of the salience network in obesity: A resting-state fMRI study: Salience Network and Obesity. Human Brain Mapping, 34(11), 2786– 2797. https://doi.org/10.1002/hbm.22104

Goulden, N., Elliott, R., Suckling, J., Williams, S. R., Deakin, J. F. W., & McKie, S. (2012). Sample Size Estimation for Comparing Parameters Using Dynamic Causal Modeling. Brain Connectivity, 2(2), 80–90. https://doi.org/10.1089/brain.2011.0057

Gutierrez, R., Fonseca, E., & Simon, S. A. (2020). The neuroscience of sugars in taste, gut-reward, feeding circuits, and obesity. Cellular and Molecular Life Sciences, 77(18), 3469–3502. https://doi.org/10.1007/s00018-020-03458-2

Han, W., Tellez, L. A., Perkins, M. H., Perez, I. O., Qu, T., Ferreira, J., Ferreira, T. L., Quinn, D., Liu, Z.-W., Gao, X.-B., Kaelberer, M. M., Bohórquez, D. V., Shammah-Lagnado, S. J., de Lartigue, G., & de Araujo, I. E. (2018). A Neural Circuit for Gut-Induced Reward. Cell, 175(3), 665-678.e23. https://doi.org/10.1016/j.cell.2018.08.049

Harding, I. H., Andrews, Z. B., Mata, F., Orlandea, S., Martínez-Zalacaín, I., Soriano-Mas, C., Stice, E., & Verdejo-Garcia, A. (2018). Brain substrates of unhealthy versus healthy food choices: Influence of homeostatic status and body mass index. International Journal of Obesity, 42(3), 448–454. https://doi.org/10.1038/ijo.2017.237

Hetherington, A. W., & Ranson, S. W. (1983). Hypothalamic Lesions and Adiposity in the Rat. Nutrition Reviews, 41(4), 124–127. https://doi.org/10.1111/j.1753-4887.1983.tb07169.x

Horst, G. J. T., & Luiten, P. G. M. (1986). The projections of the dorsomedial hypothalamic nucleus in the rat. Brain Research Bulletin, 16(2), 231–248. https://doi.org/10.1016/0361-9230(86)90038-9

Ilango, A., Kesner, A. J., Keller, K. L., Stuber, G. D., Bonci, A., & Ikemoto, S. (2014). Similar Roles of Substantia Nigra and Ventral Tegmental Dopamine Neurons in Reward and Aversion. The Journal of Neuroscience, 34(3), 817. https://doi.org/10.1523/JNEUROSCI.1703-13.2014

Jennings, J. H., Rizzi, G., Stamatakis, A. M., Ung, R. L., & Stuber, G. D. (2013). The Inhibitory Circuit Architecture of the Lateral Hypothalamus Orchestrates Feeding. Science, 341(6153), 1517–1521. https://doi.org/10.1126/science.1241812

Jennings, Joshua H., Ung, R. L., Resendez, S. L., Stamatakis, A. M., Taylor, J. G., Huang, J., Veleta, K., Kantak, P. A., Aita, M., Shilling-Scrivo, K., Ramakrishnan, C., Deisseroth, K., Otte, S., & Stuber, G. D. (2015). Visualizing Hypothalamic Network Dynamics for Appetitive and Consummatory Behaviors. Cell, 160(3), 516–527. https://doi.org/10.1016/j.cell.2014.12.026

Jung, K., Friston, K. J., Pae, C., Choi, H. H., Tak, S., Choi, Y. K., Park, B., Park, C.-A., Cheong, C., & Park, H.-J. (2018). Effective connectivity during working memory and resting states: A DCM study. NeuroImage, 169, 485–495. https://doi.org/10.1016/j.neuroimage.2017.12.067

Kühnen, P., Krude, H., & Biebermann, H. (2019). Melanocortin-4 Receptor Signalling: Importance for Weight Regulation and Obesity Treatment. Trends in Molecular Medicine, 25(2), 136–148. https://doi.org/10.1016/j.molmed.2018.12.002

Kullmann, S., Heni, M., Linder, K., Zipfel, S., Häring, H.-U., Veit, R., Fritsche, A., & Preissl, H. (2014). Resting-state functional connectivity of the human hypothalamus: Hypothalamus Functional Connectivity Networks. Human Brain Mapping, 35(12), 6088–6096. https://doi.org/10.1002/hbm.22607

Kullmann, S., Heni, M., Veit, R., Ketterer, C., Schick, F., Häring, H.-U., Fritsche, A., & Preissl, H. (2012). The obese brain: Association of body mass index and insulin sensitivity with resting state network functional connectivity. Human Brain Mapping, 33(5), 1052–1061. https://doi.org/10.1002/hbm.21268

Kwon, H. G., & Jang, S. H. (2014). Differences in neural connectivity between the substantia nigra and ventral tegmental area in the human brain. Frontiers in Human Neuroscience, 8, 41–41. PubMed. https://doi.org/10.3389/fnhum.2014.00041

Luiten, P. G., Room, P., & Lohman, A. H. (1980). Ependymal tanycytes projecting to the ventromedial hypothalamic nucleus as demonstrated by retrograde and anterograde transport of HRP. Brain Research, 193(2), 539–542. https://doi.org/10.1016/0006-8993(80)90184-5

Martín-Pérez, C., Contreras-Rodríguez, O., Vilar-López, R., & Verdejo-García, A. (2019). Hypothalamic Networks in Adolescents With Excess Weight: Stress-Related Connectivity and Associations With Emotional Eating. Journal of the American Academy of Child & Adolescent Psychiatry, 58(2), 211-220.e5. https://doi.org/10.1016/j.jaac.2018.06.039

Nieh, E. H., Vander Weele, C. M., Matthews, G. A., Presbrey, K. N., Wichmann, R., Leppla, C. A., Izadmehr, E. M., & Tye, K. M. (2016). Inhibitory Input from the Lateral Hypothalamus to the Ventral Tegmental Area Disinhibits Dopamine Neurons and Promotes Behavioral Activation. Neuron, 90(6), 1286–1298. https://doi.org/10.1016/j.neuron.2016.04.035

Palmiter, R. D. (2007). Is dopamine a physiologically relevant mediator of feeding behavior? Trends in Neurosciences, 30(8), 375–381. https://doi.org/10.1016/j.tins.2007.06.004

Park, H.-J., Friston, K. J., Pae, C., Park, B., & Razi, A. (2018). Dynamic effective connectivity in resting state fMRI. NeuroImage, 180, 594–608. https://doi.org/10.1016/j.neuroimage.2017.11.033

Penninx, B. W. J. H., & Lange, S. M. M. (2018). Metabolic syndrome in psychiatric patients: Overview, mechanisms, and implications. Dialogues in Clinical Neuroscience, 20(1), 63–73. PubMed. https://doi.org/10.31887/DCNS.2018.20.1/bpenninx

Petrovich, G. D. (2018). Lateral Hypothalamus as a Motivation-Cognition Interface in the Control of Feeding Behavior. Frontiers in Systems Neuroscience, 12, 14. https://doi.org/10.3389/fnsys.2018.00014

Preller, K. H., Razi, A., Zeidman, P., Stämpfli, P., Friston, K. J., & Vollenweider, F. X. (2019). Effective connectivity changes in LSD-induced altered states of consciousness in humans. Proceedings of the National Academy of Sciences, 116(7), 2743. https://doi.org/10.1073/pnas.1815129116

Razi, A., Kahan, J., Rees, G., & Friston, K. J. (2015). Construct validation of a DCM for resting state fMRI. NeuroImage, 106, 1–14. PubMed. https://doi.org/10.1016/j.neuroimage.2014.11.027

Rolls, E. T., Huang, C.-C., Lin, C.-P., Feng, J., & Joliot, M. (2020). Automated anatomical labelling atlas 3. NeuroImage, 206, 116189. https://doi.org/10.1016/j.neuroimage.2019.116189

Rossi, M. A., Basiri, M. L., McHenry, J. A., Kosyk, O., Otis, J. M., van den Munkhof, H. E., Bryois, J., Hübel, C., Breen, G., Guo, W., Bulik, C. M., Sullivan, P. F., & Stuber, G. D. (2019). Obesity remodels activity and transcriptional state of a lateral hypothalamic brake on feeding. Science, 364(6447), 1271–1274. https://doi.org/10.1126/science.aax1184

Rossi, M. A., & Stuber, G. D. (2018). Overlapping Brain Circuits for Homeostatic and Hedonic Feeding. Cell Metabolism, 27(1), 42–56. PubMed. https://doi.org/10.1016/j.cmet.2017.09.021

Seeley, W. W., Menon, V., Schatzberg, A. F., Keller, J., Glover, G. H., Kenna, H., Reiss, A. L., & Greicius, M. D. (2007). Dissociable Intrinsic Connectivity Networks for Salience Processing and Executive Control. Journal of Neuroscience, 27(9), 2349– 2356. https://doi.org/10.1523/JNEUROSCI.5587-06.2007

Stevens, F. L. (2011). Anterior Cingulate Cortex: Unique Role in Cognition and Emotion. J Neuropsychiatry Clin Neurosci, 6.

Stuber, G. D., & Wise, R. A. (2016). Lateral hypothalamic circuits for feeding and reward. Nature Neuroscience, 19(2), 198–205. https://doi.org/10.1038/nn.4220

Tambini, A., Ketz, N., & Davachi, L. (2009). Enhanced brain correlations during rest are related to memory for recent experiences. Neuron, 65, 280–290.

Voigt, K., Murawski, C., Speer, S., & Bode, S. (2020). Effective brain connectivity at rest is associated with choice-induced preference formation. Human Brain Mapping, n/a(n/a). https://doi.org/10.1002/hbm.24999

Zeidman, P., Jafarian, A., Seghier, M. L., Litvak, V., Cagnan, H., Price, C. J., & Friston, K. (2019). A guide to group effective connectivity analysis, part 2: Second level analysis with PEB. NeuroImage, 200, 12–25.

Zhang, S., Wang, W., Zhornitsky, S., & Li, C. R. (2018). Resting State Functional Connectivity of the Lateral and Medial Hypothalamus in Cocaine Dependence: An Exploratory Study. Frontiers in Psychiatry, 9. https://doi.org/10.3389/fpsyt.2018.00344

